# Crystal structure and molecular dynamics of human POLDIP2, a multifaceted adaptor protein in metabolism and genome stability

**DOI:** 10.1101/2020.07.24.219980

**Authors:** Anastasija A. Kulik, Klaudia K. Maruszczak, Dana C. Thomas, Naomi L. A. Nabi, Martin Carr, Richard J. Bingham, Christopher D. O. Cooper

**Author notes:** Corresponding author: Christopher Cooper.

## Abstract

Polymerase δ-interacting protein 2 (POLDIP2, PDIP38) is a multifaceted, ‘moonlighting’ protein, involved in binding protein partners from many different cellular processes, including mitochondrial metabolism, DNA replication and repair, and reactive oxygen species generation. POLDIP2 is found in multiple cellular compartments, potentially shuttled depending on its role. How POLDIP2 binds to and coordinates many different proteins is currently unknown. Towards this goal, we present the crystal structure of the ‘mitochondrial’ fragment of POLDIP2 to 2.8 Å. POLDIP2 exhibited a compact two-domain β-strand-rich globular structure, confirmed by circular dichroism and small angle X-ray scattering approaches. POLDIP2 comprised canonical DUF525 (ApaG) and YccV-like domains, but with the conserved domain linker packed tightly, resulting in an ‘extended’ YccV module. A central channel through POLDIP2 was observed which we hypothesise could influence structural changes potentially mediated by redox conditions, following observation of a modified cysteine residue in the channel. Unstructured regions were rebuilt by *ab initio* modelling to generate a model of full length POLDIP2. Molecular dynamics simulations revealed a highly dynamic N-terminal region tethered to the YccV-domain by an extended linker, potentially facilitating interactions with distal binding partners. Finally we build models of POLDIP2 interacting in complex with two of its partners in genome stability, PrimPol and PCNA. These indicate that dynamic flexibility of the POLDIP2 N-terminal and loop regions are critical to mediate protein-protein interactions.

## Introduction

Biochemical processes rarely exist in isolation in the cellular environment. Most processes comprise bridging molecules such as proteins and small organic compounds, that integrate disparate pathways to maintain cellular homeostasis. Such bridging proteins need to interact with a wide variety of partners, and how they maintain such structural plasticity is unknown. One such example is POLDIP2 (Polymerase delta-interacting protein 2, PDIP38), a poorly characterised protein involved in diverse processes including genome stability, reactive oxygen species (ROS)-mediated signaling and mitochondrial metabolism (1,2).

POLDIP2 was first identified as a partner of the p50 subunit of DNA polymerase δ (3). It is ubiquitously expressed and in addition to nuclear localisation, it is also found at cytoplasmic and mitochondrial locations, depending on cell type (reviewed elsewhere (1)). POLDIP2 is approximately 42 kDa in size, but its N-terminal targeting sequence is proteolytically cleaved to produce a ∼38 kDa protein during transit to the mitochondria (4). The N-terminal 50 residues are predicted to be mostly unstructured, followed by two predicted globular bacterial-like YccV and ApaG/ DUF525 domains of unknown function (1). POLDIP2 plays several roles in genome stability, with POLDIP2 enhancing error-free bypass of 8-oxo-G and other DNA lesions by DNA polymerases Polη, Polλ, and PrimPol (5,6). Furthermore, POLDIP2 also interacts with multiple mitochondrial components and influences mitochondrial morphology (7). POLDIP2 binds to the NADPH oxidase subunit p22^phox^, activating Nox4 production of ROS in vascular smooth muscle cells. This directly regulates cytoskeletal dynamics (8), potentially playing a role in neovascularisation and response to ischemia or other circulatory disorders. As POLDIP2 stimulates ROS production, its further role in genome stability suggests a feedback loop to protect DNA from oxidative damage (5). Furthermore, POLDIP2 knockdown protects neuronal cells from pathological Tau protein aggregation, suggesting that targeting particular POLDIP2 interactions could be beneficial in the treatment of neurodegeneration (9). POLDIP2 may also play roles in cancer, associating with proliferation-related replicative proteins (3) and also POLDIP2 knockdown suppresses lung tumour invasion (10). Conversely, POLDIP2 binds ClpX protease, downregulating lipoylation pathways and HIF-1α stabilisation, reducing cancer-related neo-angiogenesis (11).

Therefore structural determination of POLDIP2 is urgently required to understand how an otherwise small protein can ‘moonlight’, by binding and regulating diverse protein partners in processes of significant biomedical interest. Here we report the crystal structure of the POLDIP2^51-368^ (mitochondrial) fragment to 2.8 Å. We demonstrate that POLDIP2 has a rigid core structure with an extended YccV and DUF525 domains closely packed, confirmed by solution scattering and molecular dynamics simulations. We suggest that many interactions with client proteins such as PrimPol and PCNA require highly dynamic conformational changes of the unstructured N-terminal region and inter-strand loops. We also observed the presence of a central channel traversing the POLDIP2 core, and propose this could act to mediate conformational changes following the observation of a modified cysteine residue situated within.

## Results

### Origin and evolution of POLDIP2

POLDIP2 comprises an unconserved N-terminal domain (NTD), predicted to be unstructured, followed by conserved and structured YccV-like and DUF525 (ApaG) domains (Figure 1, Supplementary Figures S1 and S2), both thought to derive from bacterial ancestors (1). To assist in mapping the evolution of POLDIP2 to its structure, we performed phylogenetic analysis. Reciprocal whole genome BLAST searches found no POLDIP2 orthologues in the Fungi/Nucleariida or in eukaryotes outside the Opisthokonta (1), but did reveal a single POLDIP2 orthologue in the vast majority of holozoan clades studied, including the Filasterea and Choanoflagellatea (but notably not in the Ichythosporea) (Figure S1). Phylogenetic analyses, using Bayesian inference and maximum likelihood methodologies (Figure S2), showed that deeper internal branches recovered using both methodologies were poorly supported, however the phylogenies were consistent with the vertical inheritance of POLDIP2 since the last common ancestor of Filasterea, Choanoflagellatea and Metazoa. We therefore hypothesise that POLDIP2 originated from a fusion of the individual YccV-like and DUF525 domains during the opisthokont radiation within the Holozoa. The number of available ichthyosporean genomes is limited, but the apparent absence of the gene within the group indicates that POLDIP2 arose within Holozoa at a point after the ichthyosporean lineage split from the lineage leading to the filastereans, choanoflagellates and metazoans.

**Figure 1.**
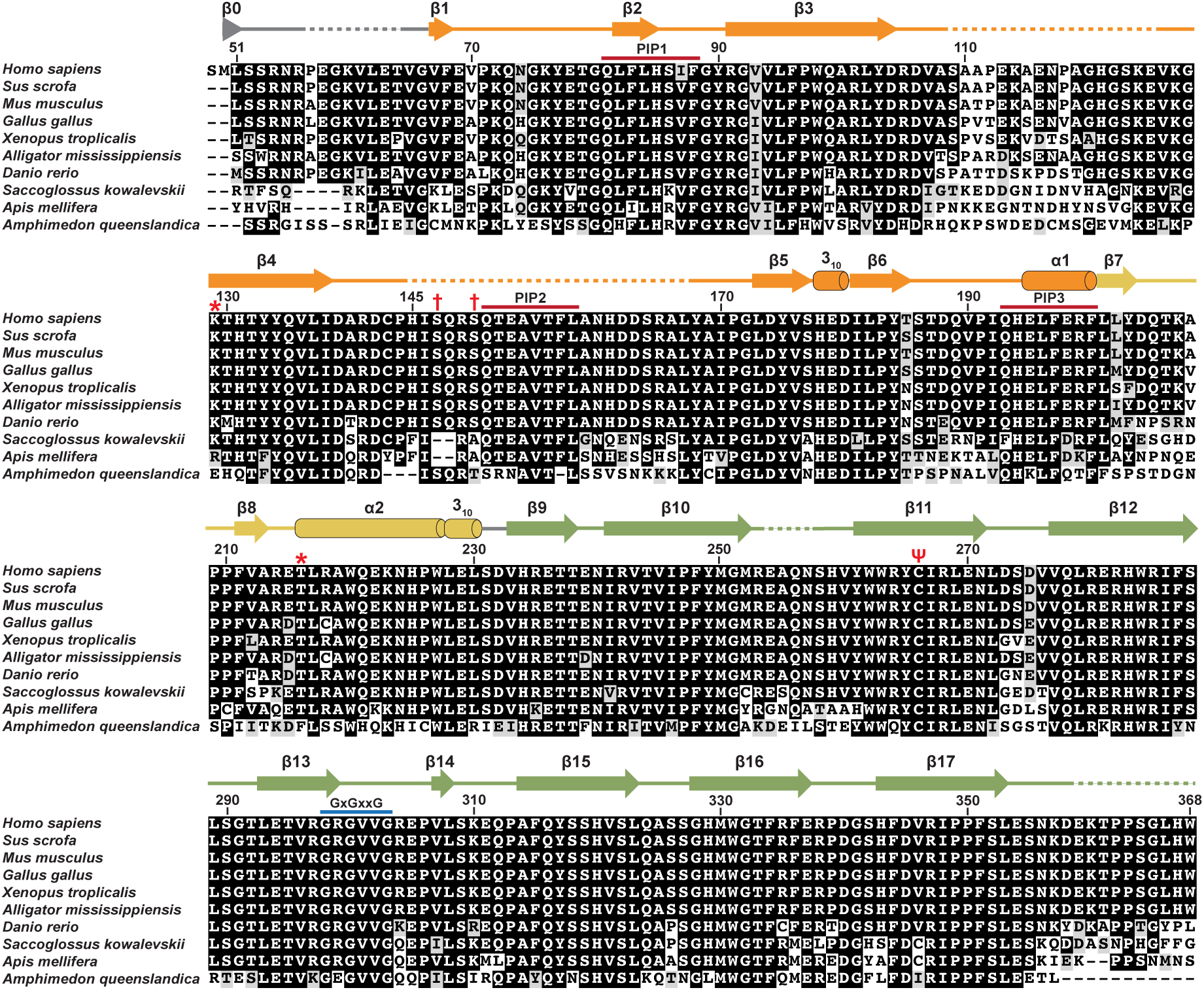
POLDIP2 structural alignment. POLDIP2 protein sequence alignment, with secondary structure assigned from the POLDIP2^51-368^ structure (PDB: 6Z9C). Alignment shading reflects amino acid identity/similarity, and secondary structure colouring reflects domain/region (orange, canonical YccV; yellow, extended YccV; green, DUF525). Residue numbering is above the alignment; Ψ represents modified Cys^266^; asterisks (*), residues crosslinking with PrimPol (6); dagger (†), ATR-phosphorylated residues (36); red lines, PCNA-interacting protein-box (3) (PIP); blue line, GxGxxG conserved motif.

### Structural determination

We initially screened a number of expression constructs in order to increase the likelihood of obtaining soluble and crystallising POLDIP2 protein (12), with POLDIP2^51-368^ expressed as an N-terminal 6xHis/thioredoxin fusion (13) being most suitable. This corresponded to the ∼38 kDa fragment resulting from post-translational removal of the mitochondrial targeting sequence, through to the intact C-terminus (Figure 1) (4). Following tag cleavage and purification, hexagonal crystals grew within 24 hours, reaching ∼0.5 mm within a week (Supplementary Figure S3a). Crystals diffracted to 2.8 Å (Supplementary Figure S3b), belonging to space group *P*6_2_ with unit cell dimensions of *a* = 120.14 Å, *b* = 120.14 Å, *c* = 49.52 Å, and α = 90.00°, β = 90.00°, γ = 120.00°. We used a molecular replacement approach to solve the POLDIP2^51-368^ structure, using bacterial HspQ (Ycc domain)(14) and human human FBxo3 (ApaG/DUF525 domain) (15). The final refined model was solved to 2.8 Å, comprising one molecule in the asymmetric unit (‘SM’ from tag, residues 51-56, 65-108, 126-144, 168-253, 258-358) with a *R*_*work*_/*R*_*free*_ factors of 0.216/0.288 respectively, 99.67% completeness and 99.6% of residues in preferred or allowed Ramachandran regions (Table 1, Supplementary Figure S3c).

**Table 1.** Data collection and structural determination statistics.

| PDB Accession Code | 6Z9C |
| --- | --- |
| **Data Collection** |  |
| Source | D8 Venture (Bruker) |
| Temperature (K) | 110 |
| Wavelength (Å) | 1.54 |
| Resolution range (Å) | 23.88–2.80 (2.95–2.80) |
| No. of measured reflections | 77725 (6274) |
| No. of unique reflections | 10246 (1488) |
| Multiplicity | 7.6 (4.2) |
| Completeness (%) | 99.7 (99.7) |
| Mean $I/\sigma(I)$ | 14.6 (1.6) |
| $R_{\text{merge}}$ | 0.139 (0.95) |
| $R_{\text{pim}}$ | 0.053 (0.516) |
| Space group | P62 |
| $a, b, c$ (Å) | 120.14, 120.14, 49.52 |
| $\alpha, \beta, \gamma$ (°) | 90.00, 90.00, 120.00 |
| **Refinement** |  |
| Resolution range (Å) | 23.88–2.80 (2.873–2.800) |
| No. of reflections | 9711 (702) |
| Completeness (%) | 99.67 (99.87) |
| $R_{\text{work}}$ | 0.216 (0.39) |
| $R_{\text{free}}$ | 0.288 (0.43) |
| $R_{\text{free}}$ test set size | 524 reflections (5.1%) |
| **Geometry** |  |
| R.m.s.d., bond lengths (Å) | 0.008 |
| R.m.s.d., bond angles (°) | 1.608 |
| Ramachandran plot |  |
| Preferred regions (%) | 228 (90.5 %) |
| Allowed regions (%) | 23 (9.1 %) |
| Outliers (%) | 1 (0.4 %) |
| **B factors (Å<sup>2</sup>)** |  |
| Mean B Value | 42.6 |
| From Wilson Plot | 43.7 |
| Total number of atoms | 2184 |

Values in parentheses represent highest resolution shell.

### Overall structure

The POLDIP2^51-368^ crystal structure exhibits a globular fold, with the YccV-like and DUF525 domains juxtaposed closely on top of each other (Figure 2a). The YccV-like domain (residues 67-200, β1-β5, 3_10_, β6, α1) comprises a 5-membered antiparallel β-sheet flanked by 3_10_ and C-terminal α-helices, reminiscent of a kinked β-barrel (orange, Figures 1 and 2a). This is very similar to the canonical YccV fold found in bacteria such as that from *E. coli* HspQ, but with extended central β3/β4 strands (3.4 Å RMSD superimposition on 5YCQ; Figure 2b, left panel) (14). An additional short β1 strand runs antiparallel to strand β6 at the edge of the β-sheet, connecting the YccV-like domain to the N-terminal region. Two long and unstructured inter-strand loops are apparent (β3-β4 and β4-β5, residues 109-125 and 145-167 respectively), owing to the lack of electron density for these regions, with the β4-β5 loop considerably longer than the equivalent loop from the HspQ YccV domain (Figure 2b, left panel). The β3-β4 loop is particularly poorly conserved in the context of the surrounding sequence (Figure 1), suggesting this region may not have a conserved role. The HspQ YccV domain is observed to form stable trimers (14), but no oligomerisation was observed for POLDIP2^51-368^ in the crystal structure, nor in solution during size exclusion chromatography (data not shown).

**Figure 2.**
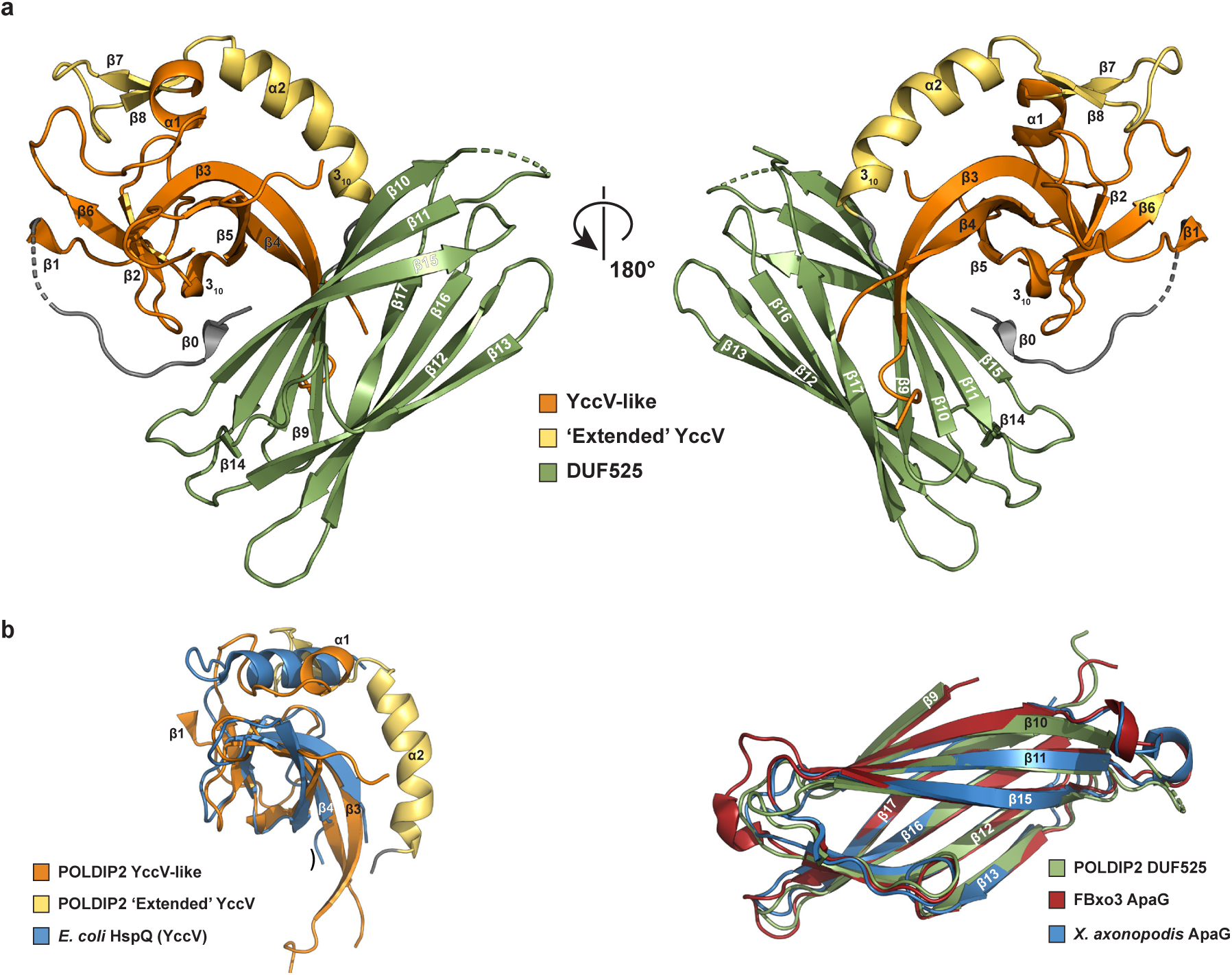
POLDIP2 overall structure. (a) Cartoon representation of the overall POLDIP2^51-368^ structure (PDB: 6Z9C). Domain colouring: orange, canonical YccV; yellow, extended YccV; green, DUF525). Cylinders, α- or 3_10_ helix; arrow, β-strand; dashed lines, disordered protein regions/missing electron density. (b) Superimposition of POLDIP2 onto structural homologues. Left panel: extended YccV domain with *E. coli* HspQ (PDB: 5YCQ; right panel: DUF525 domain with human FBx03 ApaG (PDB: 5HDW) and *Xanthomonas axonopodis* ApaG (PDB: 2F1E).

The YccV-like α1 helix leads immediately into a presumed ‘linker’ region between the YccV and DUF525 domains, comprising the α2/3_10_ helix (residues 201-230). This region is closely packed against the β3 edge of the YccV β-sheet and the α1 helix, with a significant contact area of ∼900 Å^2^ (yellow, Figure 3a). Extensive non-polar contacts are observed here, including a potential T-stacking (*π*) interaction between Phe^197^ and Phe^211^ (5.4 Å between aromatic centroids). These are capped with three hydrogen bonds between the YccV α1 helix and β7-β8 strand, and a hydrogen bond and a potential weak electrostatic interaction involving Gln^135^ on strand β4 and the α2 helix (Figure 3a). Following these extensive interactions and the high sequence conservation of the α2 helix (Figure 1), it suggests that this region is an important and structured part of the protein, rather than a simple interdomain linker. Hence, we propose this region is part of a larger ‘Extended’ YccV domain (E-YccV). Following the E-YccV a short linker (Ser^231^-Asp^232^) connects to the DUF525 domain.

**Figure 3.**
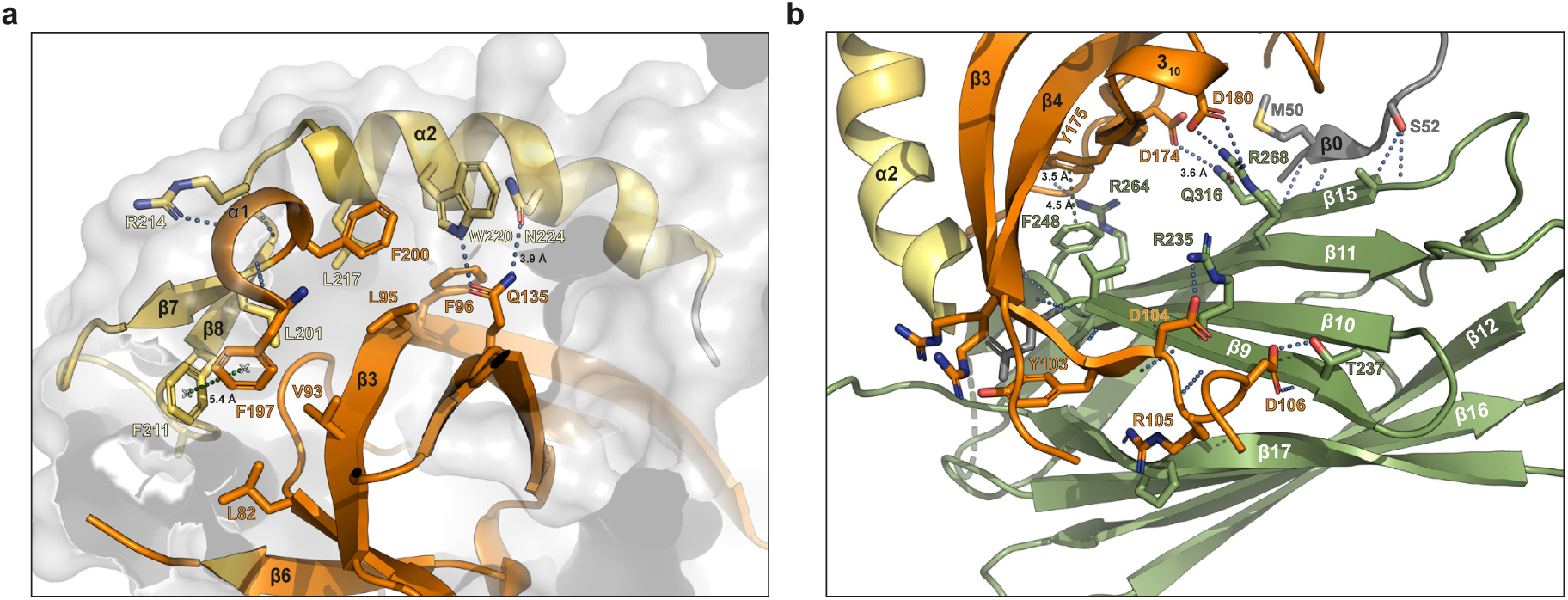
POLDIP2 domain interfaces. (a) Cartoon representation of the YccV/extended region interface. (b) Cartoon representation of the YccV/DUF525 region interface. Key residues are shown in the stick representation. Cartoon and stick colouring reflects Figure 2 (orange, YccV; yellow, extended YccV, green, DUF525). Hydrogen bonds are represented as blue dotted lines if < 3.4 Å, otherwise inter-atomic distances are labelled; green dotted lines are π-π or aromatic interactions.

The DUF525 domain (residues 233-368) exhibits a core fibronectin type III/immunoglobulin (β-sandwich) fold, comprising two opposing four-stranded antiparallel β-sheets (β9, β10, β11, β15 situated below the E-YccV domain; with β13, β12, β16, β17 opposing; Figure 2a). This is capped at either end of the fold with inter-strand loops, on one side including a short β-strand (β14), but no 3_10_ helices unlike with other DUF525 domains (Figure 2b, right panel). POLDIP2 DUF525 (residues 233-368) superimposes well onto related proteins (Figure 2b, right panel), such as human FBxo3 (5HDW; 0.7Å RMSD), the only other protein in the human genome with a DUF525 domain (15), and also bacterial ApaG homologues, such as that from *Xanthomonas axonopodis* (2F1E; 1.1Å RMSD) (16). DUF525 exhibits a significant hydrophobic core, with the β-sandwich opening up slightly to form a hydrophobic cleft between strands β13 and β15, which has been the target of drug design efforts with the equivalent region in FBxo3 (17). All loop regions in DUF525 were ordered with the exception of part of the β10-β11 loop (residues 254-257).

Part of the N-terminal loop sequence was structured in the crystal (residues 49-66), comprising a short β0 strand formed partly from Met^50^ that remained following fusion tag cleavage, with missing electron density for residues 57-64. β0 forms hydrogen bonds with the distal β15 strand of the upper antiparallel β-sheet (Figure 3b), thereby connecting DUF525 to the N-terminal region (Figure 2a).

### The POLDIP2 domain interface reveals a central channel with a modified cysteine residue

The E-YccV and DUF525 domains are closely associated, as the dipeptide linker (Ser^231^-Asp^232^) positions E-YccV directly on top of the DUF525 β-sandwich. The E-YccV/DUF525 interface involves mainly main chain and side chain hydrogen bond contacts between at least 20 residues, localised to opposite sides of the interface (Figure 3b). These are centred around interactions between the β9 strand on the edge of DUF525 and the C-terminal end of the E-YccV β3 strand and subsequent loop, and the β11 and β15 strands on the opposing DUF525 face that contacts the N-terminal β0 strand and areas around the E-YccV 3_10_ helix and β5 strand. A potential π-π stacking/aromatic interaction is found between the domains at Phe^248^ and Tyr^175^. The contact area of the E-YccV/DUF525 interface is ∼1060 Å^2^, perhaps lower than may be expected, particularly as the intra-domain E-YccV interface is ∼900 Å^2^. Surprisingly, we did not observe any residue interactions or hydrophobic-rich internal regions in the central part of the E-YccV/DUF525 interface. Rather, on closer examination we observed an internal cavity traversing the protein (Figure 4a, dashed boxes), lined with predominantly hydrophilic residues contributed from both E-YccV and DUF525 domains (Figure 4a, inset). Clear openings can be seen on opposite sides of the channel, with a narrowest channel diameter at ∼6.5 Å (Figure 4a), This channel hence accounts for the lower than expected interface contact area for E-YccV/DUF525 domains.

**Figure 4.**
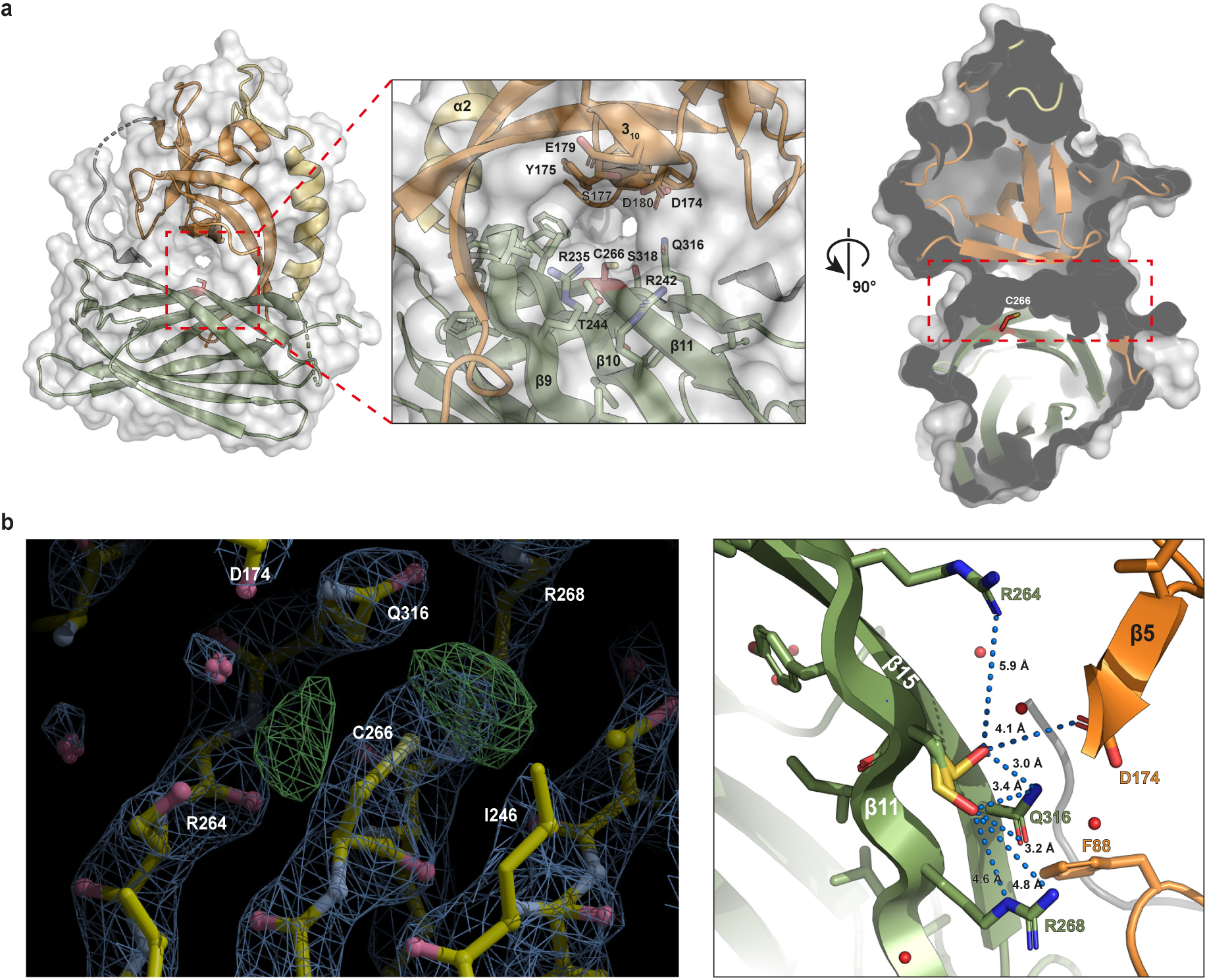
A central channel in POLDIP2. (a) Central channel traversing POLDIP2. Left panel: surface and cartoon representation, with channel marked with red dashed line. Inset: magnification of channel showing surface polar/charged residue side chains. Cys^266^ marked in stick form (red). Right panel: cross-sectional surface representation of channel, marked with red dashed line, displaying Cys^266^ in stick form (red). (b) Modification of Cys^266^ in the POLDIP2^51-368^ structure. Left panel: electron density map of POLDIP2 in the vicinity of the channel Cys^266^, with *F*_*o*_-*F*_*c*_ (green) contoured at 3.0 RMSD and 2*F*_*o*_-*F*_*c*_ (blue) contoured at 1.6 RMSD. Right panel: modelling of Cys^266^ (stick, yellow) and two potential conformations of Cys^266^ modified with sulfenic acid (stick, yellow/red), and resulting potential inter-residue distances. Key residues are shown in the stick representation. Cartoon and stick colouring reflects Figure 2 (orange, YccV; yellow, extended YccV, green, DUF525). Hydrogen bonds are represented as blue dotted lines if < 3.4 Å, otherwise inter-atomic distances are labelled; red spheres are structured water molecules.

The role of this channel is undetermined, however, we noted that Cys^266^ is situated centrally in the traversing channel, with its side chain protruding into the channel where it narrows (Figure 4a, right panel). During structural refinement we observed that Cys^266^ exhibited electron density potentially reflecting the two potential Cys^266^ rotamers that could not be accounted for by the side chain or surrounding residues (green, Figure 4b, left panel). We propose this represents a post-translational, *in crystallo* cysteine modification, as plasmid DNA sequencing and protein mass spectrometry confirmed the cysteine identity prior to crystallisation (not shown). We propose that this additional Cys^266^ density reflects an oxidised cysteine modification, and hence, its exposure to the cellular redox environment. Oxidative cysteine modifications are common in proteins, including sulfenylation (Cys-SOH) and sulfinylation (Cys-SO_2_H) (18), often modulating protein structural changes in response to ROS (19). Whilst the Cys^266^ side chain does not form hydrogen bonds with surrounding residues (Figure 4b, right panel), we modelled sulfenic acid into the two potential rotameric positions, finding that the Cys^266^-SOH could potentially participate in hydrogen bonding with Gln^316^ and weaker electrostatic interactions elsewhere, both within and between domains. Although sulfenic acid does not satisfy the additional electron density, it demonstrates that potential Cys^266^ oxidative modification could potentially change the local non-covalent bonding properties and interactions with other residues, thereby influencing structural or dynamic alterations, affecting downstream interactions with partner proteins. Cys^266^ is conserved in the great majority of species examined across the Holozoa (Figure 1 (Ψ); Supplementary Figure S1), also supporting a conserved function. Furthermore, POLDIP2 is known to regulate cellular redox potential by upregulating Nox4 activity to produce ROS (8). Hence, the existing link of POLDIP2 to redox control suggests the possibility that the exposed Cys^266^ in the traversing channel could act as a redox sensor, with modifications (oxidative or otherwise) modulating POLDIP2 interactions.

### POLDIP2^FL^ exhibits a rigid core structure, with significant conformational disorder in termini and loops

In order to further validate the POLDIP2 crystal structure, we also performed biophysical analysis on the protein in solution. The high β-strand content of POLDIP2^51-368^ was confirmed with circular dichroism analysis (Figure 5a). Furthermore, although we hypothesised that the channel that runs between the E-YccV and DUF525 domains may have a functional role, it remains a possibility that this channel in the domain interface is an artefact of crystal packing. Therefore we performed small angle X-ray scattering (SAXS, Figure 5b) to investigate the solution structure of POLDIP2^51-368^. The SAXS Guinier plot was linear at low *q* values consistent with a monodisperse system (Figure 5c), and the Kratky plot (Figure 5d) is consistent with a globular/partially unfolded structure (20), suggesting some contribution from the long, disordered loop regions. The *P*(r) distribution indicates a single globular unit (Figure 5e), rather than well-separated subunits displaying multiple maxima (21). The Porod volume of 56100 nm^3^ and the real/reciprocal space *R*_g_ values (22.49/22.72 Å) are consistent with the dimensions and molecular weight of monomeric POLDIP2^51-368^. Taken together, the SAXS data are entirely consistent with the asymmetric unit seen in the crystal structure with E-YccV associating tightly with DUF525 as a bound unit in solution. Furthermore, the crystallographic B factors (Figure 6a) indicate a generally rigid core E-YccV/DUF525 with limited motion, with only intermediate motion likely for the inter-strand loop regions and the YccV extension α2/β7-β8 strands.

**Figure 5.**
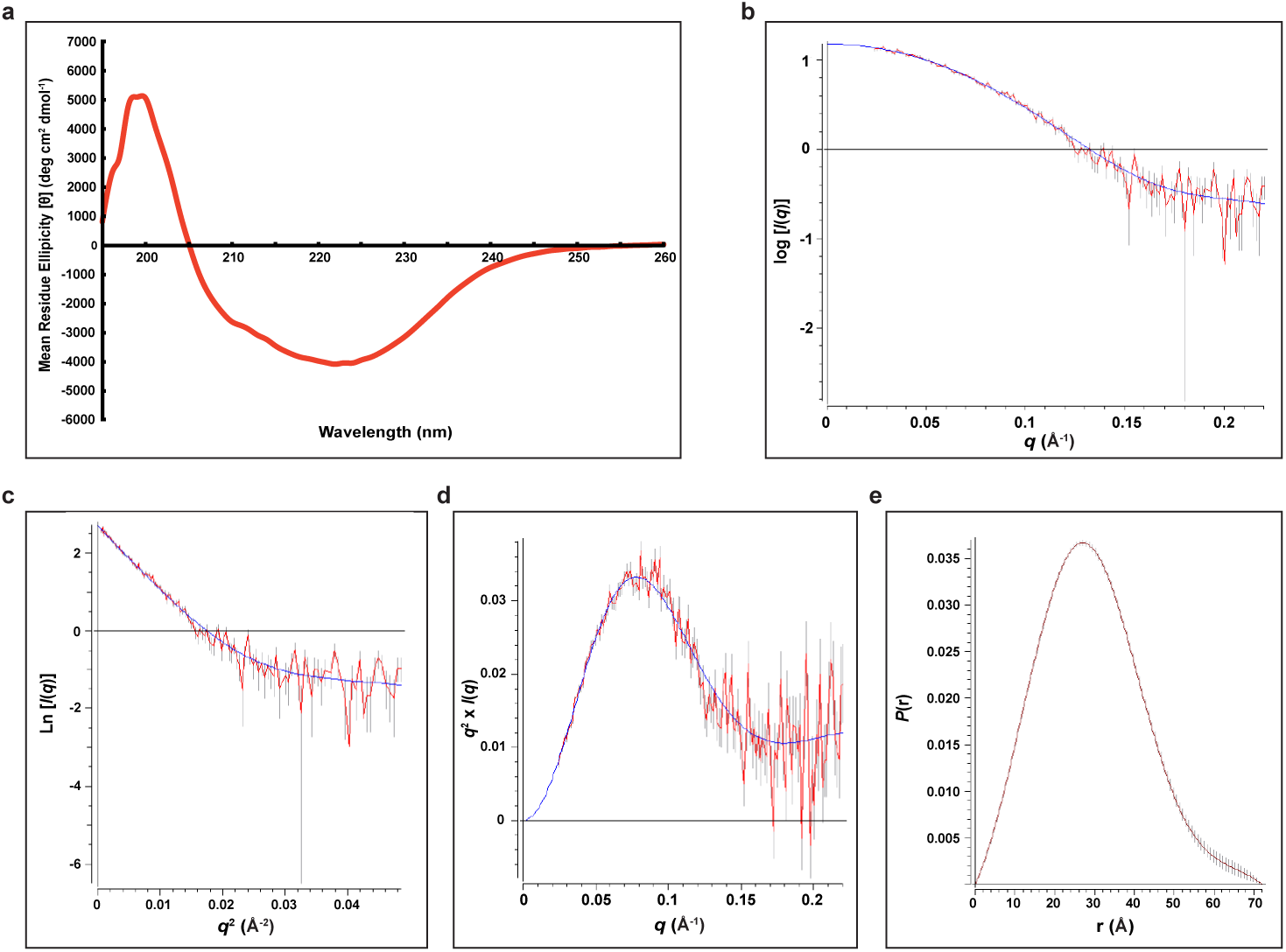
Circular dichroism and SAXS solution structural analysis of POLDIP2^51-368^. (a) Far-UV circular dichroism analysis of POLDIP2^51-368^, normalised to buffer blank. (b) raw SAXS scattering data of POLDIP2^51-368^, following averaging and buffer subtraction in PRIMUS, plotted as log *I*/*q*. (c) Guiner analysis of SAXS data from (b), noting the linear shape at low *q*^2^ values. (d) Kratky plot (*q*^2^ x *I*(*q*) versus *q*) of SAXS data from (b). (e) Distance distribution function *P*(*r*) representation of SAXS data from (b), using a D_max_ of 72.22. For SAXS analyses: red lines, raw data; blue lines, fitted curves; grey lines, error bars.

**Figure 6.**
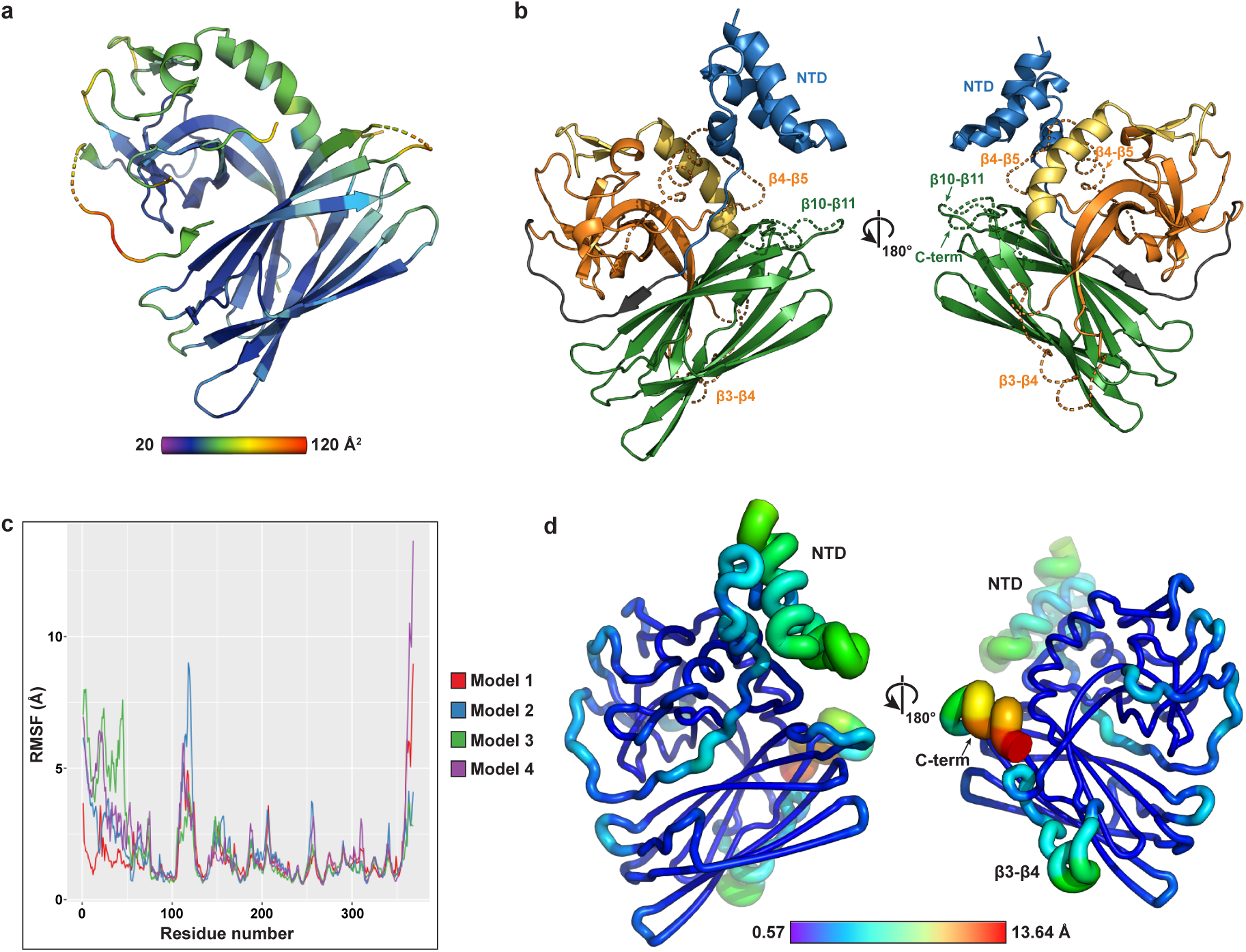
Structural modelling and dynamics of POLDIP2. (a) Temperature (B) factors of POLDIP2^51-368^ plotted on cartoon representation. (b) Representative Robetta model of POLDIP2^FL^ (#4). Domain colouring as in Figures 1 and 2a, excepting N-terminal domain in blue cartoon representation (NTD) and modelled loops as dashed lines. (c) RMSF analysis of all four full-length Robetta models. Root mean square fluctuation (RMSF, deviation of atoms with respect to reference structure) plotted against residue position. (d) RMSF analysis of 100 ns molecular dynamics simulation of POLDIP2^FL^ model 4, positioned with respect to structures in Figure 2a. Colour scale and chain thickness varies from blue (low fluctuation) to red (high fluctuation).

In addition to the N-terminal 50 residues excluded from the POLDIP2^51-368^ expression construct, several external loops are missing from the crystal structure following conformational disorder (Figure 1). These include several protein-protein interaction sites, such as the NTD contacting PrimPol (6) and a PCNA-interacting protein (PIP) box in loop β4-β5 (3). In order to gain further understanding of full-length POLDIP2 (POLDIP2^FL^), we rebuilt these missing loop regions and the NTD using Robetta (22), with POLDIP2^51-368^ as a template structure for comparative modelling. Five models were generated, each of which differed in the orientation of loops and the NTD (Supplementary Figure S4). The NTD is predicted to form a globular helical bundle, the precise arrangement of which varies between the models, presumably reflecting genuine conformational heterogeneity (Figure 6b; Supplementary Figure S4b). However, the NTD of model 5 was observed to pass through a loop of the E-YccV domain, forming a knot topology and therefore was not included in further analysis (not shown). Loops β3-β4, β4-β5 and the C-terminus are predicted to form short α-helices, the lengths and orientations of each differs between the four models. Residues Leu^51^-Gly^66^ form an extended loop, connecting the NTD to the β1 strand of the E-YccV domain (grey, Figure 6b). The crystal structure resolves part of this linker occupying a narrow groove between the E-YccV and DUF525 domains, forming an β0 strand against the DUF525 β15 strand (Figure 2a), potentially stabilising the interaction between the two domains.

To gain further insight into possible conformational changes and how these may influence functional protein-protein interactions, the four POLDIP2^FL^ models were subjected to molecular dynamics simulations. The overall drift from the initial model was evaluated by calculating C_α_ RMSD values over 100 ns simulation (Supplementary Figure S5a). All four models diverged significantly during the initial stages, and models 1-3 reached a plateau indicating a stable average conformation, while model 4 continued to diverge indicating convergence had not occurred (Supplementary Figure S5a). Longer 1000 ns simulations indicate that the N-terminal α-helix is particularly mobile (Supplementary Figure S5b). Structural displacements were measured by RMSF analysis, revealing regions of stability and local flexibility (Figures 6c and 6d, Supplementary Figure S4). Conformational flexibility within the simulation was consistent with B-factor analysis (and missing residues) from the crystal structure (Figure 6a). All four models revealed high mobility for both loop β3-β4 and the C-terminal residues 358-368 (Figure 6c). The NTD had high mobility during simulations of models 2-4, whereas in model 1 the NTD had lower mobility, forming a more stable interaction with the E-YccV and DUF525 domains (Figure 6c, Supplementary Figure S4). Although models 1-3 all converged to a stable conformation over the simulation time, the configuration of the NTD remained quite different between them. Furthermore, during the simulation of model 4 the NTD had high mobility with concomitant high RMSF values (Figure 6c and 6d). Taken together, this data shows that the NTD has high mobility, tethered to the E-YccV domain by a long linker, stabilised by a short β-strand situated between the E-YccV and DUF525 domains. It is therefore possible that a conformational change altering the E-YccV/DUF525 domain interface may destabilise this long linker and free the NTD to form interactions at a distance.

### POLDIP2 surface structural features and interaction motifs

POLDIP2 is observed to interact with a great number of protein partners from across multiple processes, particularly those from mitochondrial metabolism and genome stability (1). Direct binding or biochemical modulation is observed with eight DNA polymerases alone (2), but how POLDIP2 exhibits such plasticity in binding is unknown. Although the POLDIP2^FL^ model presented here suggests the NTD could be structured, its highly dynamic nature and heterogeneity of length and sequence composition across the Holozoa (Supplementary Figure S1) are reminiscent of characteristics seen in intrinsically disordered proteins (23). This could potentially facilitate binding to many different partners, relying on short linear motifs (24) such as the N-terminal/mitochondrial-targeting helix critical for binding PrimPol (6), or highly charged regions (23). The surface of the POLDIP2^51-368^ exhibits highly positively charged and basic regions on either face (Figure 7a), with the POLDIP2^FL^ model surface demonstrating a contribution of significant additional positive charge resulting in a polarised surface (Figure 7b), as the reverse face containing the β0 strand is less charged overall in POLDIP2^FL^ models. This could facilitate more generalised electrostatic interactions with a wide variety of negatively charged partners, somewhat explaining the promiscuity of POLDIP2 interactions. The POLDIP2^51-368^ surface also has localised patches of hydrophobicity on both faces of the structure (Figure 7b) particularly around the DUF525 cleft previously described, potentially contributing to protein-protein interaction interfaces.

**Figure 7.**
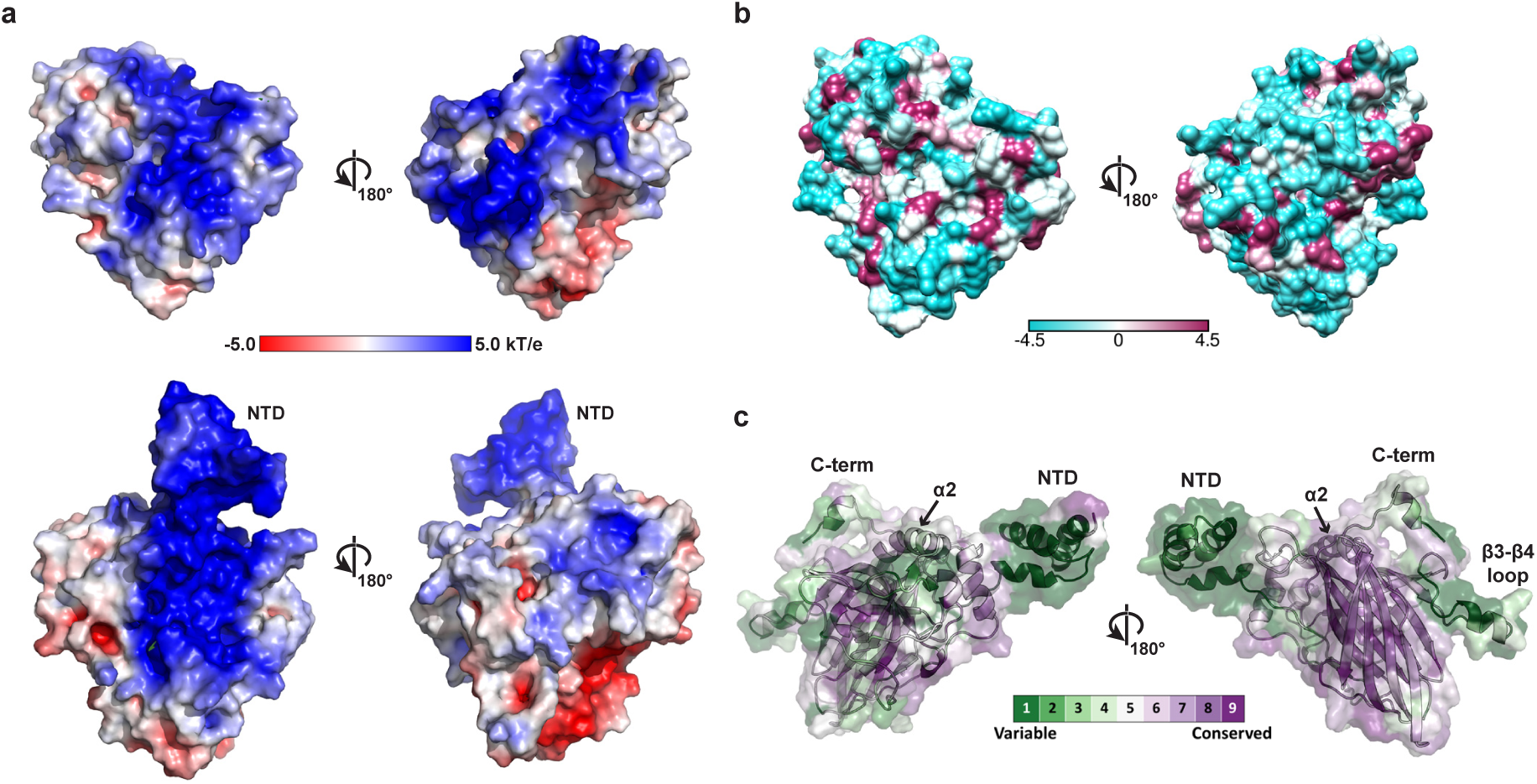
Conserved and surface features of POLDIP2. (a) POLDIP2 electrostatic surface, calculated in ABPS (47), orientated with respect to structures in Figure 2a. Upper panel: POLDIP2^51-368^ crystal structure; lower panel full-length POLDIP2 model 4. NTD, N-terminal domain. (b) POLDIP2^51-368^ surface hydrophobicity, orientated with respect to structures in Figure 2a. Scale bar represents Kyte and Doolittle scale (69). (c) evolutionary conservation calculated with ConSurf was mapped onto the POLDIP2^FL^ model 4 (purple, most conserved; green, most variable).

We also mapped protein sequence evolutionary conservation onto the POLDIP2^FL^ model surface (Figure 7c). The exposed DUF525 domain β-sandwich is particularly conserved (Figure 7c, right model), with the β16-β17 loop in this DUF525 region directing interaction with the CEACAM1 cell-cell adhesion receptor, binding to POLDIP2 to regulate its subcellular localisation between the cytoplasm and nucleus (25). The significant evolutionary and hence structural conservation around this POLDIP2 region (Figure 1, 7c) suggests a functional relationship with CEACAM1 in POLDIP2 subcellular trafficking may occur across all vertebrates. The most variable regions were the dynamic N- and C-termini, unstructured β3-β4 loop and the E-YccV domain β3-β4 antiparallel β-strand, suggesting that flexibility and/or sequence composition rather than sequence could be more important in directing interactions with some POLDIP2 binding partners. The DUF525 β13 strand has a surface-exposed β-strand and is therefore accessible for interaction with other proteins by β-strand addition. Sitting at the base of the domain it forms crystal contacts comprising an intermolecular antiparallel β-sheet with the equivalent β13 strand on an adjacent monomer (not shown). POLDIP2 also contains a conserved glycine-rich motif associated with pyrophosphate or nucleotide binding (GxGxxG, Figure 1) in the DUF525 region (raspberry, Figure 8a), (26), situated beside an arginine-rich cluster (27). The GxGxxG motif was not observed to bind nucleotides in DUF525 or ApaG homologues (15,16). However, both the glycine and arginine-rich regions are important for the stimulation of PrimPol DNA synthesis, with the arginine-rich cluster also stimulating dNTP binding (27).

**Figure 8.**
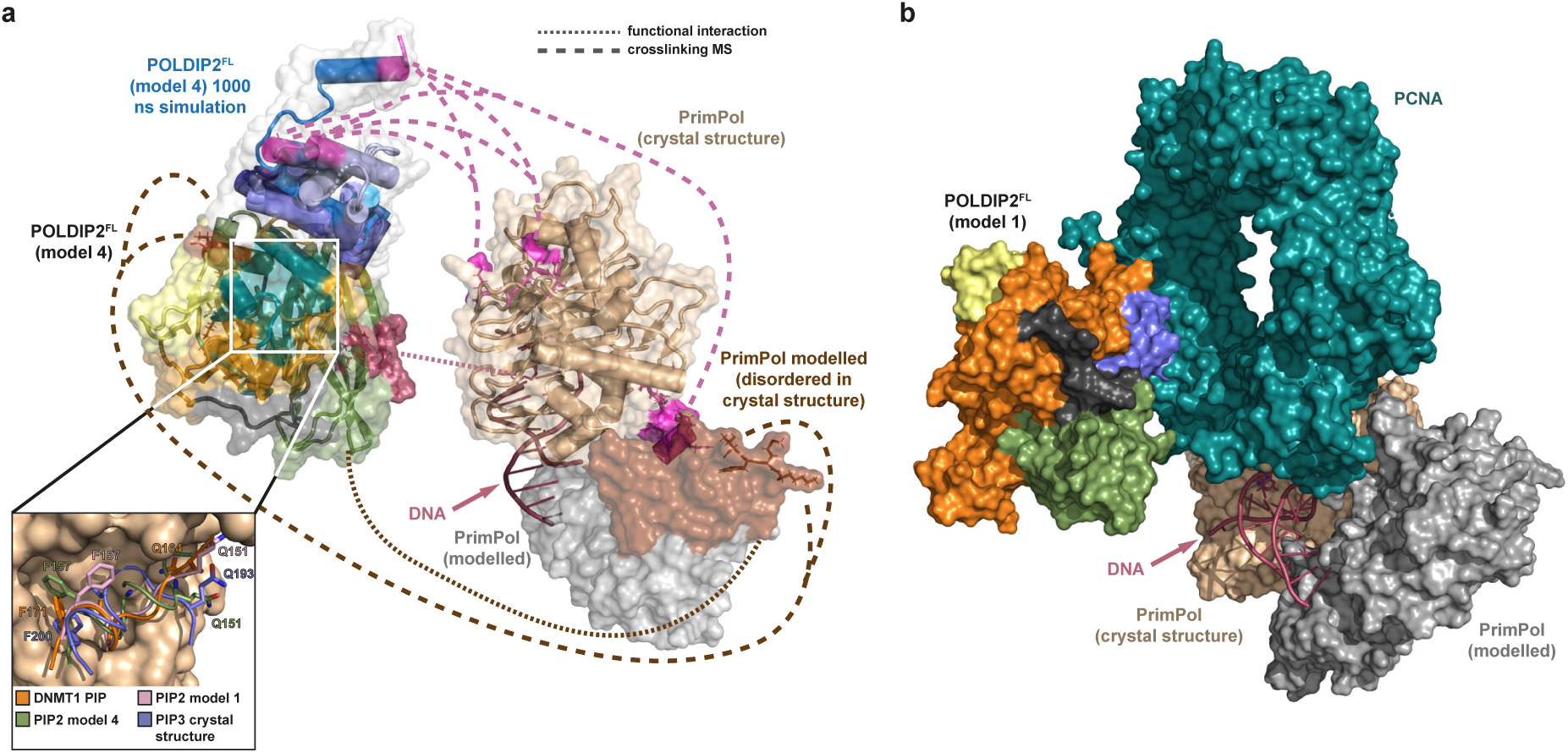
Structural modelling of POLDIP2^FL^-PrimPol^FL^ protein complex. (a) Structural model of POLDIP2^FL^ interactions with PrimPol^FL^. Molecules are separated to illustrate interactions, with matching colours and connecting lines representing functional (raspberry, dotted lines) (27) or crosslinking mass spectrometric (pink/brown, dashed lines) studies (6). Protein structures were derived using Robetta from the POLDIP2^51-368^ crystal structure determined herein (6Z9C), and the PrimPol^1-354^ crystal structure (5L2X). POLDIP2^FL^ model 4 (Figure 6) was used as a representative model, with 1000 ns simulation (Supplementary Figure S5b) timepoints superimposed for the NTD to illustrate temporal heterogeneity. Dark green/teal, potential PIP boxes in POLDIP2^FL^ model 4. Inset: PIP boxes from the POLDIP2^51-368^ crystal structure and POLDIP2^FL^ models 1 and 4, superimposed onto the DNMT1 PIP box bound to human PCNA (63KA), with consensus residues represented as sticks (39). (b) Structural model of human PCNA (63KA) abrogating POLDIP2^FL^-PrimPol^FL^ interactions by steric occlusion, with POLDIP2 and PrimPol separated for illustrative purposes. The human PCNA-DNMT1 PIP-peptide complex (teal green) was superimposed against the POLDIP2^FL^ model 1 PIP2 box, with POLDIP2^FL^-domain colouring as in Figures 1 and 2a. PrimPol is positioned according to (a) but distanced, to demonstrate mutual exclusivity of PCNA/PrimPol positioning when juxtaposed to POLDIP2.

## Discussion

### Structural insights into POLDIP2 interactions with PrimPol

POLDIP2 interacts directly with a number of proteins involved in DNA replication and repair, particularly DNA polymerases (reviewed elsewhere (2)). In particular, POLDIP2^FL^ stimulates DNA binding, DNA synthesis and processivity during 8-oxoguanine translesion bypass for PrimPol (an archaeo-eukaryotic primase (28)), Polλ and Polη DNA polymerases (5,6). It is also observed to interact with Pol *ζ* (via REV7) and REV1 (29), Polβ (30), and also to partially stimulate Pol*ι* (5). How POLDIP2 achieves binding across structurally analogous but evolutionarily divergent families of DNA polymerases (31) is currently unknown. To gain further insight, we derived a model of POLDIP2-PrimPol interactions (Figure 8a), consistent with both the POLDIP2 crystal structure and modelling presented here, and previous reported experimental observations. We manually arranged the POLDIP2^FL^ and PrimPol^FL^ protomers (based on predictive Robetta models), using known contact points from bissulfosuccinimidyl suberate (BS3) crosslinking mass spectrometry (XlMS) (6) and functional inference from biochemical analyses (27).

Following the requirement of the glycine/arginine-rich region for PrimPol stimulation described (27), the region was orientated in our model close to where the incoming dNTP binds to the primer/template (p/t) DNA in PrimPol (Figure 8a). POLDIP2 was further directionally orientated to PrimPol using XiMS contact points (6). Here, the N-terminal helix of the POLDIP2 NTD contacts multiple spatially separated regions around the PrimPol catalytic site (pink, Figure 8a) in keeping with our model, following the highly dynamic nature of the NTD seen both between different POLDIP2 models and simulations (Figure 8a, Supplementary Figures S4 and S5). One such binding site however is on the far distal side of PrimPol (residues 343-352). However, an interaction is still feasible from our model, as the modelled POLDIP2 NTD could potentially fully unfold and stretch to reach this distal face of PrimPol, particularly if the constraining β0 strand that connects the NTD in a β-sheet to the DUF525 domain was to become destabilised, granting the NTD significantly greater conformational flexibility. POLDIP2 NTD binding to PrimPol may be only partially specific, following the multiple PrimPol sites bound by the NTD N-terminal helix (6). Following the very basic nature of this helix from its conserved arginine residues (Supplementary Figure S1), this suggests POLDIP2 interactions may involve generalised features such as electrostatic charge, other than structured or conserved structural elements. The spatial arrangement of the complex protomers however distanced the modelled PrimPol disordered loop (residues 201-260; brown, Figure 8a) from its corresponding crosslinked POLDIP2 residues in the E-YccV domain (Lys^129^ and Thr^216^, Figure 1, asterisks), to opposite distal faces of the complex. It should be noted however that our representation of PrimPol^FL^ is a single predictive model, and it is feasible that the disordered PrimPol loop could stretch across to interact with the distal face of POLDIP2. This PrimPol loop also functionally interacts with C-terminal/DUF525 region (27), which is closer than the E-YccV (crosslinking) contacts in our model. This suggests that significant conformational flexibility and dynamic motion is required from both POLDIP2 and PrimPol partners, to reconcile reported functional and crosslinking features. An alternative explanation for POLDIP2 binding to multiple distant sites on PrimPol however, could result from multiple POLDIP2 molecules bound to a single PrimPol unit, as positively cooperative binding kinetics were previously observed for the POLDIP2-mediated stimulation of PrimPol (6).

From our model, we speculate that the POLDIP2 glycine/arginine-rich region could potentially provide additional active site residues in *trans* to facilitate dNTP binding close to the PrimPol p/t DNA region, with POLDIP2 potentially also assisting PrimPol DNA binding following the strong positive charge in this region of the DUF525 domain (Figure 7a) enclosing the p/t DNA, analogous to DNA polymerase processivity factors (32). This requires the POLDIP2 NTD however, as although C-terminal fragments alone (residues 217-368) can stimulate PrimPol DNA synthesis, full stimulation of DNA synthesis and DNA binding by PrimPol is only observed for POLDIP2^FL^ (6,27), suggesting cooperativity of binding or catalysis. Furthermore, POLDIP2 is unlikely to directly tether DNA polymerases to the DNA, as although the YccV domain is associated with hemi-methylated DNA binding in bacteria (33), POLDIP2 was not previously seen to bind directly to DNA directly (5,6). Recent reports however indicate POLDIP2 can directly bind to p/t DNA structures, but the full POLDIP2 molecule including the NTD is required (27), suggesting some contextual DNA sequence or structural specificity. A likely role therefore of the POLDIP2 NTD, is to clamp and stabilise a potential cooperative ternary complex between POLDIP2, p/t DNA and PrimPol, potentially helping to enclose the p/t DNA with its positive charge, in order to stimulate DNA synthesis by potentiating DNA-PrimPol binding. Further structural studies on POLDIP2-PrimPol complexes are required to test this hypothesis. It is tempting to speculate that such a mechanism could be the basis for POLDIP2-mediated stimulation of Polδ, Polλ and Polη. However, as these are *bone fide* DNA polymerases from the divergent DNA polymerase families B, X and Y respectively, and not an archaeo-eukaryotic primase like PrimPol, further biochemical and structural studies are required.

### POLDIP2 interactions with PCNA may be dependent on conformational flexibility

POLDIP2 was originally identified as binding partner of p50, a subunit of the replicative DNA polymerase δ (Polδ), and also its processivity factor proliferating cell nuclear antigen (PCNA) (3). POLDIP2 stimulates Polδ DNA synthesis and also PCNA binding, potentially by bridging the molecules, with the POLDIP2 NTD required for full stimulation (4). PCNA plays a major role when replicative DNA polymerases such as Polδ encounter DNA damage that impedes the replication fork. This induces PCNA monoubiquitylation by the RAD18–RAD6 ubiquitin ligase, which in turn recruits generally Y-family translesion DNA polymerases to bypass the damage (34). Polη is one such DNA polymerase recruited by ubiquitylated PCNA, with Polη interacting via its UBZ ubiquitin-binding domain (35), notably with which it also interacts with POLDIP2 (29). As POLDIP2 regulates Polη nuclear localisation in unstressed cells by sequestering it in the cytoplasm (36), but, as POLDIP2 also binds to PCNA, a greater understanding of the relationship of POLDIP2 with PCNA is crucial to understand their interplay.

POLDIP2 contains three putative PIP box sequences (PIPs) situated in the E-YccV domain, with PIP1 and PIP3 ordered in the crystal structure, but PIP2 present in the disordered β4-β5 loop (Figure 1) (3). None of the PIPs exactly match the canonical QXXΨXXF (F/Y) motif, where Ψ is an aliphatic residue, as all only exhibit a single aromatic residue (37), (although it is recognised that broader ‘PIP-like’ motifs are still able to bind to PCNA (37,38)). Mapping the PIPs onto the POLDIP2^51-368^ crystal structure and POLDIP2^FL^ models juxtaposes PIP1 and PIP3 into close proximity (teal green, Figure 8a), with PIP2 also located nearby in some models, but demonstrating positional heterogeneity between POLDIP2^FL^ models following conformational flexibility of the β4-β5 loop. PIP1 is unlikely to interact with PCNA however, as it is comprised of strand β2, yet, most PIP boxes exhibit an α- or 3_10_ helix (37,38). Moreover, PIP1 is situated away from the protein surface, hence sterically occluded from binding PCNA. We superimposed PIP3 from the POLDIP2^51-368^ crystal structure (comprising the α1 helix) and PIP2 from POLDIP2^FL^ models (containing a predicted α-helix), to the DNMT1 PIP box (6K3A) (39), which also lacks one aromatic residue from the canonical PIP motif. The PIP2 and PIP3 α-helices superimpose well to the DNMT1 PIP 3_10_ helix, with the PIP2/3 terminal phenylalanine orientated towards the PCNA hydrophobic pocket (or able to as an alternative rotamer), and the N-terminal PIP glutamine residues are situated around, but not within the PCNA Q pocket (Figure 8a, inset). PIP3 phenylalanine and glutamine residue side chains point towards the POLDIP2 core rather than to the protein surface however, therefore PIP3 would need to re-orientate 180° around the helical axis in order for PCNA to bind, although the crystal structure’s B factors and modelling RMSF values of this region suggest such a motion is unlikely (Figure 6).

POLDIP2^FL^ model RMSF values and the lack of electron density in the crystal structure indicate that PIP2 is present on a conformationally flexible loop (Figure 6c, Supplementary Figure S4). Moreover, POLDIP2^FL^ model 1 presents PIP2 as pointing away from the core POLDIP2, which would allow PCNA to access PIP2. We modelled PCNA bound to PIP2 in POLDIP2^FL^ model 1 using the PCNA-DNMT1 superimposition described (Figure 8b). This clearly shows PCNA being able to bind to the POLDIP2 surface, and although part of the NTD is trapped within the superimposition, the NTD conformational heterogeneity could allow PCNA to bind here unimpeded. When PCNA binding to POLDIP2 is combined with our model of the POLDIP2-PrimPol complex (Figure 8b), it can be clearly seen that PCNA would sterically occlude PrimPol from interacting with POLDIP2, preventing PCNA access to both the NTD and glycine/arginine-rich DUF525 region. PCNA is not known to interact with PrimPol, although addition of PCNA to both PrimPol and POLDIP2 results in PrimPol inhibition (6). This is consistent with our model, as not only would PCNA interacting at PIP2 block POLDIP2 from stimulating PrimPol, POLDIP2 could stabilise PCNA on p/t DNA structures to inhibit PrimPol, as previously suggested (6). Here, the positively charged NTD could potentially stabilise the POLDIP2-PCNA complex, following its requirement of the NTD for DNA binding on some p/t templates (27). Although PrimPol is required for replication fork progression under both normal and DNA-damaging conditions (40), the mutual exclusivity of PCNA/PrimPol binding to PCNA could help prevent aberrant PrimPol recruitment to the replication fork when PCNA is in complex with Polδ. POLDIP2 has been reported to be phosphorylated on Ser^147^ and Ser^150^ by the ATR kinase following UV irradiation (36). These two serine residues are found immediately beside PIP2 (Figure 1) and with phosphorylation adding bulky negatively charged groups, the binding properties of PIP2 would likely be changed. Hence, future studies should address if post-translational modification of POLDIP2 can influence the interaction with PCNA.

### POLDIP2 is a redox sensor and modulator protein?

The surprising identification of the hydrophilic channel between the E-YccV and DUF525 domains led us to hypothesise about its function (Figure 4a). The observation of additional electron density around the Cys^266^ side chain present in the channel (Figure 4b) prompted us to speculate that this could reflect an oxidised post-translational modification of the Cys^266^ (or modification by covalent binding by an as yet unidentified small molecule ligand), given that it is exposed to the cellular environment via the channel. POLDIP2 is involved in a number of cellular processes relating to redox status, including generation of ROS by stimulating Nox4 (8), and POLDIP2 is downregulated under hypoxic conditions which stabilises HIF-1α resulting in significant metabolic changes (11). Furthermore, POLDIP2 facilitates translesion bypass across oxidised DNA lesions resulting from ROS damage (5). Therefore an ability to sense redox state via Cys^266^ modification could potentially result in POLDIP2 structural changes, as such modifications could change local residue side chain interactions or hydrogen bonding patterns between residues surrounding the Cys^266^, as we observed when sulfenic acid was modelled at this position (Figure 4b). These could result in wider conformational or dynamic changes, including effects on the interdomain interface given the location of Cys^266^ between the E-YccV and DUF525 domains (Figure 4b). Some domain communication is likely, given that the E-YccV domain is reported to inhibit DUF525-mediated stimulation of PrimPol when the NTD is missing in POLDIP2^51-368^ (27). Such regulation has been observed in oxidative regulation of Src kinase activity (19), and is also seen in many other types of proteins (18,41).

Large conformational changes are not expected in the core POLDIP2 structure, following examination of structure B factors and modelling simulations (Figure 6). However, small conformational changes could potentially destabilise the weakly hydrogen bonded β0-strand at the edge of the upper DUF525 β-sandwich (Figure 1b), allowing even greater dynamic flexibility of the upstream NTD. Such conformational flexibility of the NTD is likely to be key to many POLDIP2 protein interactions (Figure 8a). Release of the β0-strand could also potentially promote opening of the E-YccV/DUF525 interface, following the propensity of internal hydrophilic residues (which would become externalised on opening the channel). However, our dynamics simulations, structure B factors, and in-solution SAXS analysis instead support a tight single globular unit (Figure 5), rather than a tandem domain arrangement. It is possible of course that gross structural rearrangements could occur on POLDIP2 binding to a partner, therefore structural studies on POLDIP2 complexes will be important.

### Conclusions

We present the crystal structure of POLDIP2, demonstrating that the core protein is a β-strand rich globular protein, comprising an evolutionarily conserved extended (E-)YccV domain, juxtaposed on top of a DUF525 β-sandwich domain. Evolutionary analysis supports fusion of these domains occurring soon after the establishment of the Holozoa during eukaryotic evolution. We demonstrate that POLDIP2 is a stable and rigid molecule at its core, with modelling and simulations suggesting dynamic flexibility of the N-terminal domain and loop regions being important in mediating protein interactions, inferred from modelling POLDIP2 associations with PCNA and PrimPol, key proteins in human DNA replication and repair. We also noted a channel traversing the POLDIP2 core, which we speculate could house a conformational switch, owing to the observed modification of Cys^266^ contained within. Whilst we anticipate any conformational changes to be slight, these could still have dramatic effects on the disordered loop and terminal regions on POLDIP2. Future structural determination of POLDIP2 in complex with its interaction partners will be vital to test these hypotheses, towards determining how POLDIP2 can act as a central nexus, connecting redox metabolism and genome stability.

## Methods

### Cloning and recombinant protein expression and purification

Human POLDIP2^51-368^ was amplified by PCR from plasmid pETM33 encoding FL-POLDIP2 (a kind gift from Professor B. van Loon), using Phusion DNA polymerase (New England Biolabs, Ipswich, MA, USA) and primers POLDIP2-f001 (5’-TACTTCCAATCCATGCTCTCGTCCCGAAA CCGAC-3’) and POLDIP2-r000 (5’-TATCCACCTTTACTGTCACCAGTGAAGGCCTGAGGG-3’). Purified PCR products were cloned by a ligation independent approach into the pNH-TrxT expression vector (GenBank GU269914.1, encoding an N-terminal TEV-cleavable 6xHis-thioredoxin fusion tag) (13). Plasmid constructs were confirmed by DNA sequencing and transformed into *Escherichia coli* BL21 (DE3) Rosetta2™ (Novagen^®^). Bacterial cells were cultured at 37°C in Terrific Broth containing 50 μg/ml kanamycin in Ultra Yield™ baffled flasks (Generon, Slough, UK). Recombinant POLDIP2^51-368^ expression was induced at OD_600_ of 2.5 by the addition of isopropyl-β-D-1-thiogalactopyranoside to 0.1 mM and cells incubated overnight at 18°C, prior to harvesting and storage of cell pellets at -80°C.

Thawed cells were resuspended in buffer A (50 mM HEPES (pH 7.5), 5% (v/v) glycerol, 500 mM NaCl, 10 mM imidazole, 4 mM β-mercaptoethanol, 0.5 mg/ml hen egg white lysozyme, 5U/ml Basemuncher nuclease (Abcam, Cambridge, UK) and protease inhibitors (1 mM phenylmethylsulphonyl fluoride, 1 mM benzamidine-HCl)), and disrupted by sonication on ice. Lysates were clarified by centrifugation (20000 x g, 30 min, 4°C) an applied to a 2 ml Ni-NTA immobilised metal affinity chromatography (IMAC) gravity flow column (QIAGEN, Hilden, Germany). Columns were washed in 10 column volumes (CV) of buffer A, followed by 10 CV wash buffer (buffer A with 30 mM imidazole), and eluted in 5x 2 ml fractions of buffer A containing 300 mM imidazole. Fractions were analysed by SDS-PAGE and relevant fractions pooled and cleaved overnight with 1:20 mass ratio of 6xHis-tagged TEV protease, with concurrent dialysis in buffer B (20 mM HEPES (pH 7.5), 5% (v/v) glycerol, 500 mM NaCl, 10 mM imidazole, 4 mM β-mercaptoethanol). TEV protease and additional contaminants were removed with a repeated Ni-NTA IMAC column, and flow through and relevant elutions by a stepwise 20 mM imidazole elution in buffer B. Pooled fractions were finally separated by size exclusion using a HiLoad 16/600 Superdex S200 column (GE Healthcare, Chicago, IL, USA) in buffer B, but with β-mercaptoethanol substituted with 1 mM dithiothreitol. Protein concentration was calculated from OD_280_ using molecular mass and extinction coefficients, and LC/ESI-TOF mass spectrometry used to confirm protein identity. Proteins were concentrated using 10 kDa MWCO centrifugal concentrators (VIVAproducts, Littleton, MA, USA).

### Crystallization and structural determination

Purified POLDIP2^51-368^ protein was concentrated to 20 mg/ml and sparse matrix crystal screens were performed at 293K, using a ratio of 1:1 with mother liquor in 2 μl sitting drops. Crystals were obtained in 0.2 M calcium acetate, 0.1 M sodium cacodylate, 40% PEG300, pH 6.5 and flash-cooled directly in liquid nitrogen without addition of further cryoprotectant. X-ray diffraction data were collected using a Bruker D8 Venture source coupled with a CMOS-PHOTON II detector (Bruker, Billerica, MA, USA). Data reduction was performed using Proteum3 (Bruker) and SCALA (Evans, 2006). Structure solution was by molecular replacement in Phaser (McCoy, 2007) using models based on 5HDW (ApaG/DUF525 domain) and 5YCQ (YccV domain). Search models were first modified by deletion of surface loops and removal of side chains by PDBSET (CCP4, Winn et al 2004,). Density modification was conducted using DM before automatic chain tracing in BUCCANEER (Cowtan 2006). Structure refinement then proceeded by alternating between real-space refinement in *Coot* (Emsley and Cowtan, 2010) and reciprocal space refinement in REFMAC5. Data collection and refinement statistics are shown in Table 1, with Ramachandran analysis performed in RAMPAGE (42)). Secondary structure was assigned using STRIDE (43) and PDBsum (44). Structure visualisation was performed in PyMOL or Chimera (45), interfaces determined in PISA (46), and electrostatic calculations were performed with APBS (47).

### Molecular dynamics simulations

The N-terminal region of POLDIP2 (residues 1-50) and several external loops are missing from the POLDIP2^51-368^ crystal structure. These were rebuilt using the crystal structure as a template by comparative modelling in Robetta (22), generating five models. A full-length PrimPol model was generated in Robetta using the PrimPol^1-354^ crystal structure (5L2X) (48). The POLDIP2 models were used to initiate five separate 100 ns simulations by molecular dynamics using the Gromacs package (49) with CHARMM27 force field. Each model was solvated in a dodecahedron box with a minimum distance to edge of 1.0 nm using the SPC/E water model with 0.1 M NaCl and then neutralized by addition of additional NaCl ions. Structures were minimized by steepest descent until the maximum force was < 1000 kJ/mol/nm. The system was then heated to 300 K using NVT conditions followed by stabilization of pressure under NPT conditions. Long-range electrostatic interactions were calculated using Particle Mesh Ewald method with 1 nm cutoff. Production runs were either 100ns or 1000 ns with Berendsen temperature coupling and Parrinello-Rahman pressure coupling. Trajectories were analysed using the Bio3D package in R (50).

### Small angle X-ray scattering

***(SAXS)*** Scattering data were acquired for purified POLDIP2^51-368^ using a Bruker Nanostar instrument. Protein samples were buffer matched (20 mM HEPES (pH 7.5), 5% (v/v) glycerol, 500 mM NaCl) by overnight dialysis at 4°C. Scattering data was collected from 100 μl of 3.4 mg/ml POLDIP2^51-368^ in a 1.5 mm bore quartz glass capillary, under vacuum for 10 exposures of 1800 seconds. Data were averaged and buffer scattering was subtracted using PRIMUS in the ATSAS package (51). *R*_g_ values were calculated using the Guiner approximation in PRIMUS.

### Circular dichroism spectroscopy

Circular dichroism (CD) experiments were performed on a Chirascan spectrophotometer Applied Photophysics, Beverly, USA). Far-UV spectra (180-260 nm) were obtained in a 1 mm pathlength quartz cuvette (Hellma, Plainview, USA), with 0.2 ml of 0.45 mg/ml protein (in 50 mM sodium phosphate, pH 7.5) at 5°C. The average of four spectra were taken with 1 nm resolution, 1 second response time, 2 nm bandwidth, with buffer blank spectra taken under identical conditions and subtracted from the data. Data were analysed with the DICHROWEB software suite (52).

### Sequence and phylogenetic analysis

Sequence manipulation was performed in Geneious PRIME (https://www.geneious.com). 2020.1.2 POLDIP2 orthologues were found using a reciprocal homology approach (53) using BLASTP (54) with default parameters. *Homo sapiens* POLDIP2 (NP_056399.1) was used as a query sequence against specific genomes in the NCBI GenBank database (55), or the *Mnemiopsis* genome project BLAST server (56). Domain homology searching used the SMART (57) and PFAM (58) databases. Protein sequence alignment was in MAFFT v7.450 (59) with default parameters implemented in Geneious, and visualised in Jalview (60). Ambiguously aligned regions and gaps were removed with Gblocks (61). A Bayesian inference phylogeny was created in MrBayes 3.2.6 (62) through the Cipres Science Gateway server (63). The analysis was run with a mixed amino acid model and a four category gamma distribution to correct for among site rate variation. The MCMC analyses consisted of 5,000,000 generations using two parallel chain sets, run at default temperatures. The sampling frequency was 1,000, with a burnin value of 1,250. A maximum likelihood phylogeny was created with raxmlGUI 2.0 (64), using the thorough bootstrapping methodology with 1,000 bootstrap replicates. The ML tree was generated from 100 starting parsimony trees, using the PROTCAT model and the JTT amino acid substitution matrix (65), which was recovered in the MrBayes analysis as the most appropriate matrix. Empirical amino acid frequencies were used in the ML analysis. Phylogenetic trees were rendered in FigTree v1.3.1(66). Residue evolutionary conservation was calculated with ConSurf (67) using the PDB for Robetta model 4 and protein sequence alignments from MAFFT, followed by rendering in PyMOL.

## Supporting information

Full supplementary information

## Data availability

The structure presented in this paper was deposited in the Protein Data Bank (PDB) under the code 6Z9C.

## Acknowledgements and additional comments

The authors thank Professor Barbara van Loon (Norwegian University of Science and Technology) for the kind gift of plasmid pETM33 encoding POLDIP2^FL^, and the University of Huddersfield for funding. We thank Dr. Ed Bolt for constructive reading of this manuscript. The authors declare that they have no conflicts of interest with the contents of this article. During preparation of this manuscript, the authors were aware of a related preprint submission (68).

## Author Contributions

CDOC conceived and supervised the study; AAK, MC, RJB and CDOC designed experiments; AAK, KM and CDOC provided new tools and reagents; AAK, KM, NLAN, RJB and CDOC performed experiments; RJB performed molecular dynamics studies; MC, DCT and CDOC performed bioinformatics and phylogenetic analyses; RJB and CDOC solved and analysed protein structures; AAK, RJB and CDOC performed and analysed SAXS; RJB, MC and CDOC wrote and revised the manuscript.

## Abbreviations

CD: circular dichroism;
dNTP: deoxyribonucleotide phosphate;
FL: full-length;
NTD: N-terminal domain;
PCNA: proliferating cell nuclear antigen;
PIP: PCNA interacting protein;
p/t DNA: primer-template DNA;
POLDIP2: Polymerase δ-interacting protein 2;
POLDIP2^FL^: full-length POLDIP2;
PrimPol^FL^: full-length PrimPol;
RMSD: root mean square deviation;
RMSF: root mean square fluctuation;
ROS: reactive oxygen species;
SAXS: small angle X-ray scattering;
XlMS: crosslinking mass spectrometry.

