## Supplementary material for "Crystal structure and molecular dynamics of human POLDIP2, a multifaceted adaptor protein in metabolism and genome stability": Full supplementary information

[2] Current address: Astbury Centre for Structural Molecular Biology, School of Molecular and Cellular Biology, Faculty of Biological Science, University of Leeds, Leeds, LS2 9JT

[3]Current address: Interfaculty Institute of Biochemistry, University of Tübingen, 72074, Tübingen, Germany.

List of contents:

**Supplementary Figure S1. Protein sequence alignment of POLDIP2 orthologues from major eukaryotic groups, elucidated from reciprocal BLASTP searches.**

**Supplementary Figure S2. Bayesian inference phylogeny of POLDIP2 protein sequences.**

**Supplementary Figure S3. Crystallisation and data collection of POLDIP2$^{51\text{-}368}$.**

**Supplementary Figure S4. Structural representation and molecular dynamics simulations for all full-length POLDIP2 models.**

**Supplementary Figure S5. POLDIP2$^{FL}$ molecular dynamics time simulations.**

**Supplementary Figures**

### FIGURE S1

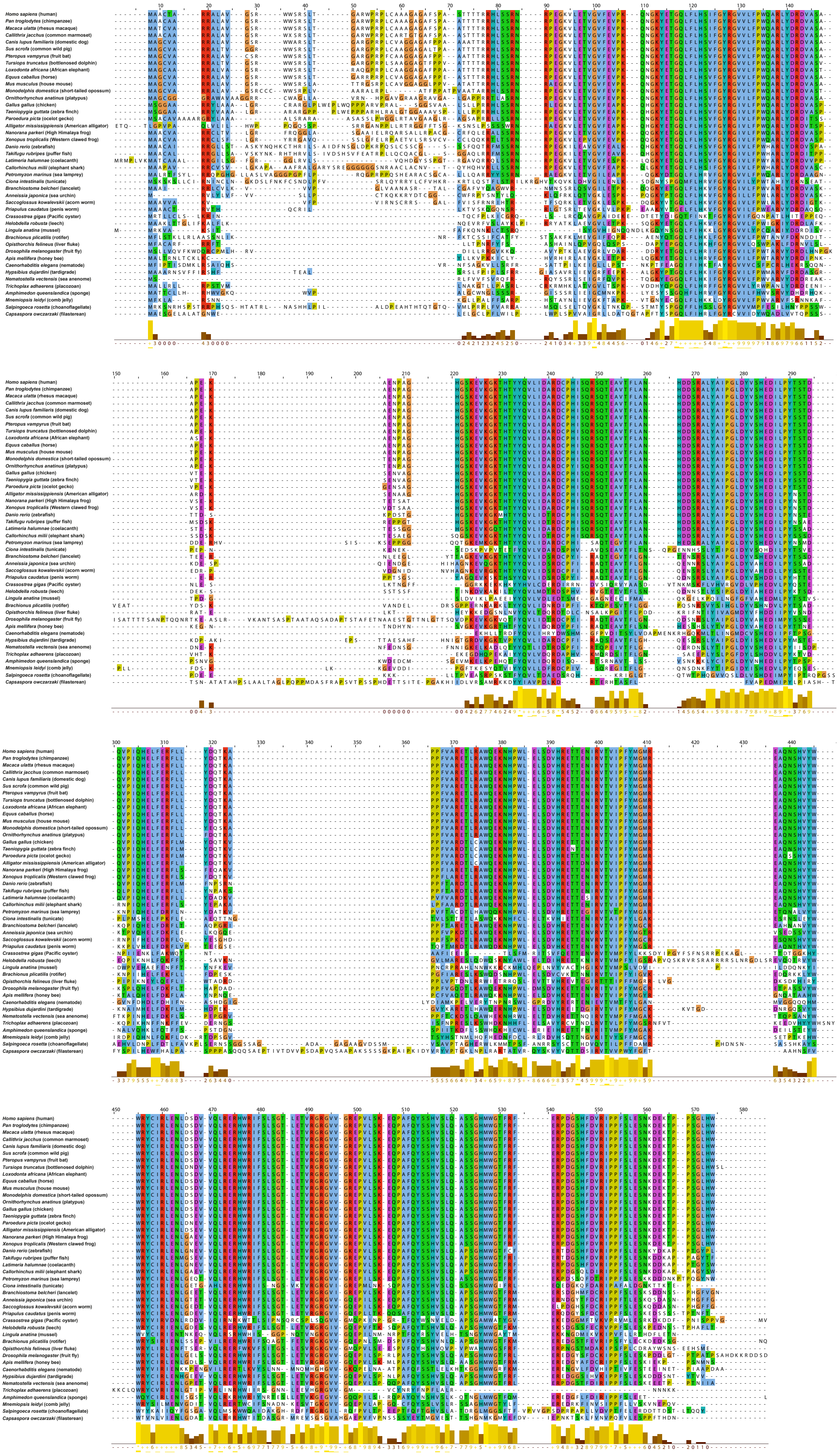
Homo sapiens (human)
Pan troglodytes (chimpanzee)
Macaca ulatta (rhesus macaque)
Callithrix jacchus (common marmoset)
Canis lupus familiaris (domestic dog)
Sus scrofa (common wild pig)
Pteropus vampyrus (fruit bat)
Tursiops truncatus (bottlenosed dolphin)
Loxodonta africana (African elephant)
Equus caballus (horse)
Mus musculus (house mouse)
Monodelphis domestica (short-tailed opossum)
Ornithorhynchus anatinus (platypus)
Gallus gallus (chicken)
Taeniopygia guttata (zebra finch)
Paroedura picta (ocelot gecko)
Alligator mississippiensis (American alligator)
Nanorana parkeri (High Himalaya frog)
Xenopus tropicalis (Western clawed frog)
Danio rerio (zebrafish)
Takifugu rubripes (puffer fish)
Latimeria halumnae (coelacanth)
Callorhinchus milii (elephant shark)
Petromyzon marinus (sea lamprey)
Ciona intestinalis (tunicate)
Branchiostoma belcheri (lancelet)
Anneissia japonica (sea urchin)
Saccoglossus kowalevskii (acorn worm)
Priapulus caudatus (penis worm)
Crassostrea gigas (Pacific oyster)
Helobdella robusta (leech)
Lingula anatina (mussel)
Brachionus plicatilis (rotifer)
Opisthorchis felineus (liver fluke)
Drosophila melanogaster (fruit fly)
Apis mellifera (honey bee)
Caenorhabditis elegans (nematode)
Hypsibius dujardini (tardigrade)
Nematostella vectensis (sea anemone)
Trichoplax adhaerens (placozoan)
Amphimedon queenslandica (sponge)
Mnemiopsis leidyi (comb jelly)
Salpingoeca rosetta (choanoflagellate)
Capsaspora owczarzaki (filasterean)

**Supplementary Figure S1. Protein sequence alignment of POLDIP2 orthologues from major eukaryotic groups, elucidated from reciprocal BLASTP searches.** Alignment was generated using MAFFT in Geneious PRIME and visualised in Jalview. Residue position is above the alignment, residue colouring is according to ClustalX, and panel below the alignment is Jalview conservation score. Sequence and GenBank identifiers were: *Homo sapiens* (NP_056399.1), *Pan troglodytes* (XP_016787681.1), *Macaca ulatta* (NP_001248642.1), *Callithrix jacchus* (XP_008995455.2), *Canis lupus familiaris* (NP_001240832.1), *Sus scrofa* (XP_003358222.2), *Pteropus vampyrus* (XP_011358283.1), *Tursiops truncatus* (XP_019782617.1), *Loxodonta africana* (XP_003416901.1), *Equus caballus* (XP_023508851.1), *Mus musculus* (NP_080665.1), *Monodelphis domestica* (XP_001368593.1), *Ornithorhynchus anatinus* (XP_028938338.1), *Gallus gallus* (NP_001304285.1), *Taeniopygia guttata* (NP_001232073.1), *Paroedura picta* (GCF45383.1), *Alligator mississippiensis* (XP_019345287.1), *Nanorana parkeri* (XP_018412666.1), *Xenopus tropicalis* (NP_001017098.1), *Danio rerio* (NP_997879.1), *Takifugu rubripes* (XP_003968776.1), *Latimeria halumnae* (XP_006011053.1), *Callorhinchus milii* (XP_007894669.1), *Petromyzon marinus* (XP_032823499.1), *Ciona intestinalis* (XP_002121208.2), *Branchiostoma belcheri* (XP_019641134.1), *Anneissia japonica* (XP_033123282.1), *Saccoglossus kowalevskii* (XP_006824194.1), *Priapulus caudatus* (XP_014665443.1), *Crassostrea gigas* (XP_011415219.2), *Helobdella robusta* (XP_009026644.1), *Lingula anatina* (XP_013400176.1), *Brachionus plicatilis* (RNA06801.1), *Opisthorchis felineus* (TGZ54899.1), *Drosophila melanogaster* (NP_649540.1), *Apis mellifera* (XP_006559902.2), *Caenorhabditis elegans* (NP_498703.2), *Hypsibius dujardini* (OQV20687.1), *Nematostella vectensis* (XP_032226378.1), *Trichoplax adhaerens* (XP_002114156.1), *Amphimedon queenslandica* (XP_019848720.1), *Mnemiopsis leidyi* (AGCP01016403), *Salpingoeca rosetta* (XP_004988627.1), *Capsaspora owczarzaki* (XP_004349208.2).

**FIGURE S2**

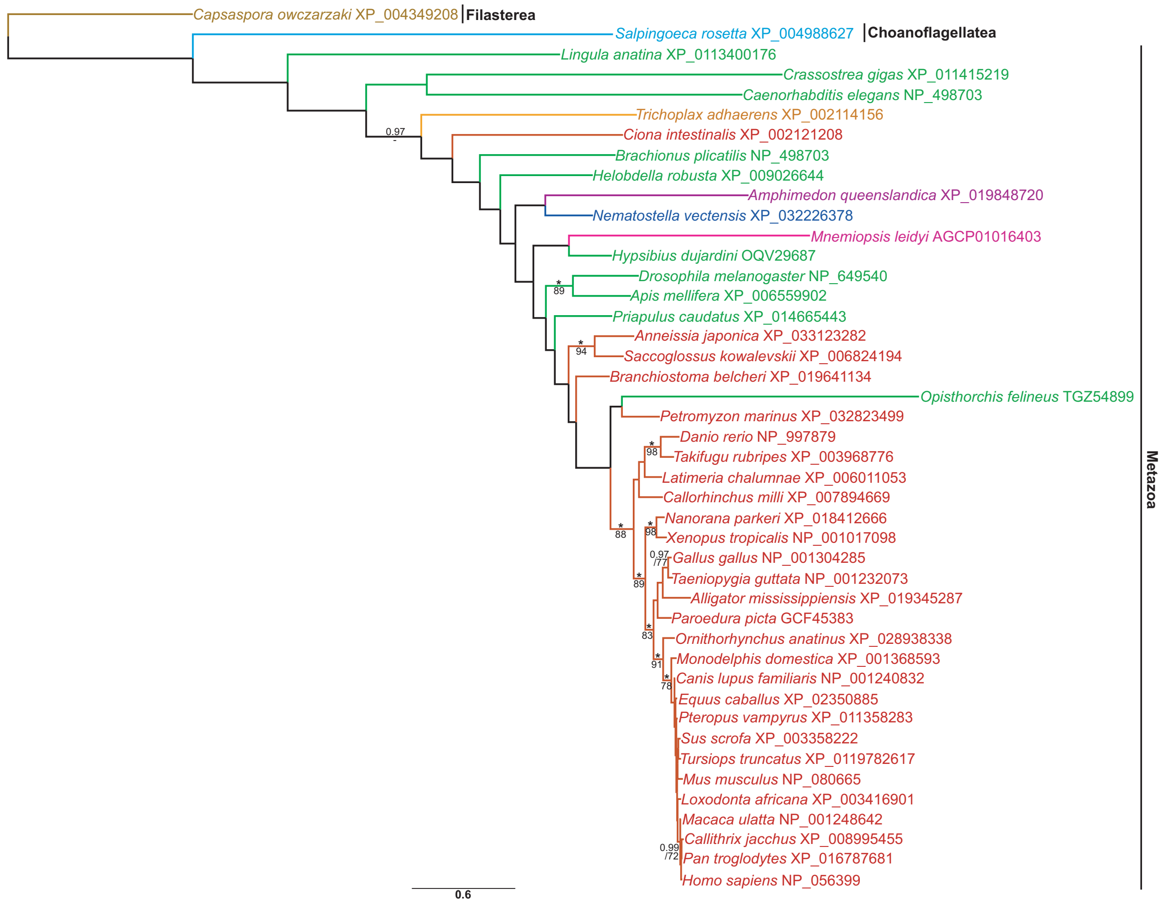


**Supplementary Figure S2. Bayesian inference phylogeny of POLDIP2 protein sequences.** The phylogeny was constructed from 368 aligned amino acid positions using the PROTCAT model, with the JTT substitution matrix, and estimated amino acid frequencies. Values for biPP and mlBP are shown above and below the branches respectively. 1.00 biPP and 100% mlBP are both denoted by "*". Values <70% mlBP and <0.97 biPP are denoted by "-". The scale bar represents the number of substitutions per site. Colour code: Brown: Filasterea; Light Blue: Choanoflagellatea; Purple: Porifera; Pink: Ctenophora; Orange: Placozoa; Dark Blue: Cnidaria; Green: Protostomia; Red: Deuterostomia.

**FIGURE S3**

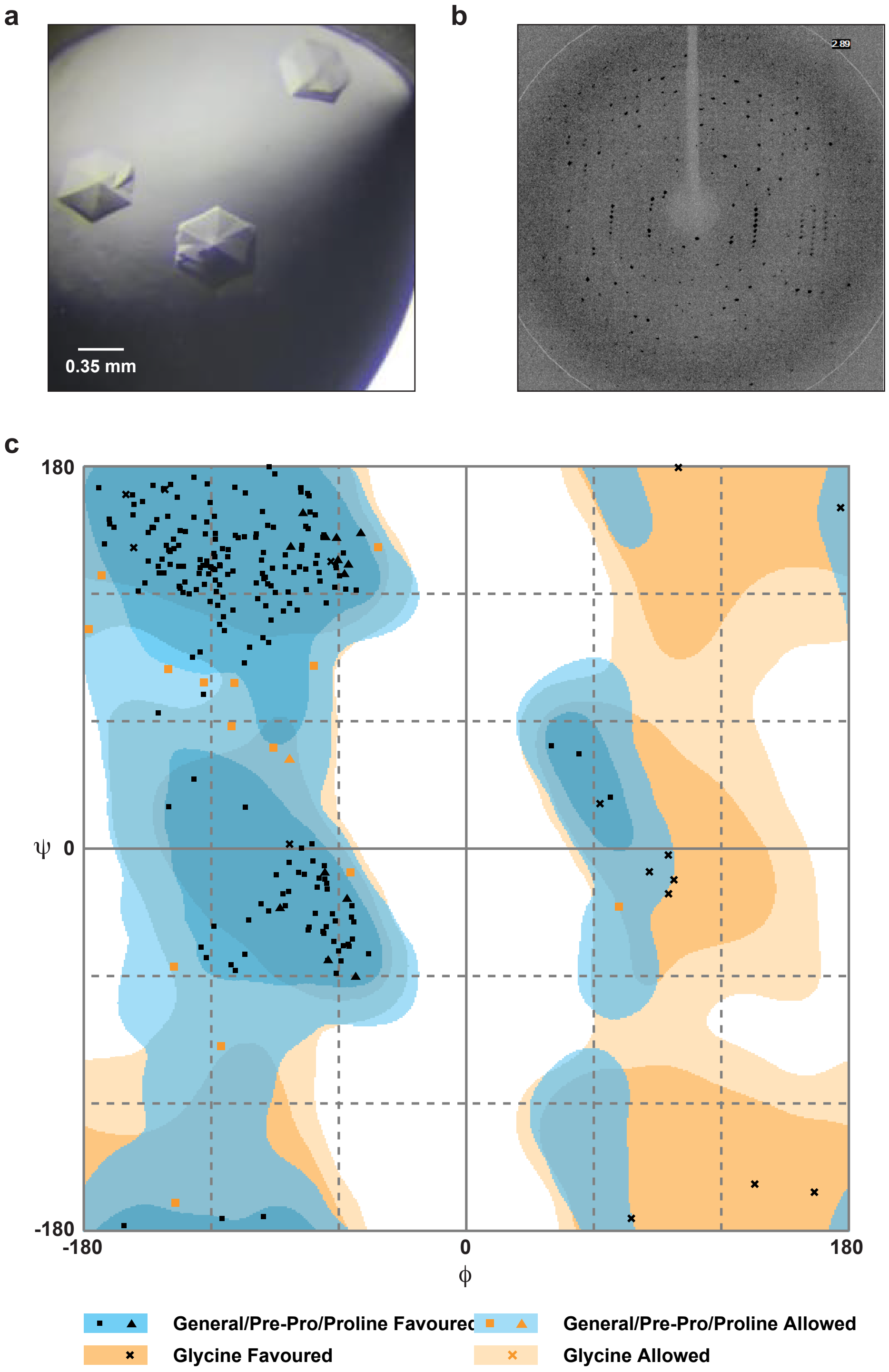


**Supplementary Figure S3. Crystallisation and data collection of POLDIP2$^{51\text{-}368}$.** (a) protein crystals of POLDIP2$^{51\text{-}368}$, appearing after 1 week. (b) X-ray diffraction pattern of POLDIP2$^{51\text{-}368}$ crystal. (c) Ramachandran plot of the ψ/ϕ main chain angles from the solved structure of POLDIP2$^{51\text{-}368}$ (PDB: 6Z9C), with 99.6% of residues in preferred or allowed regions.

**FIGURE S4**

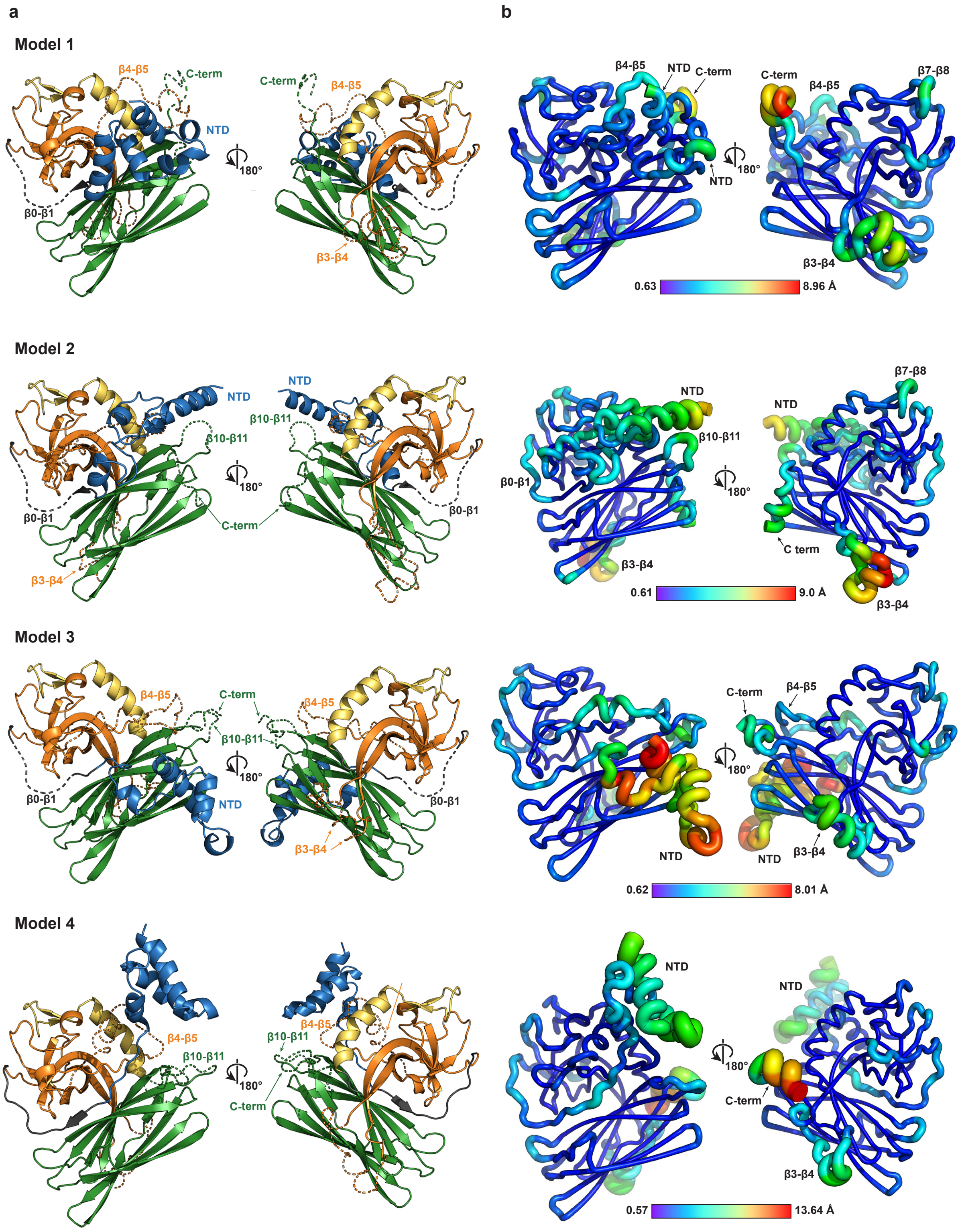


**Supplementary Figure S4. Structural representation and molecular dynamics simulations for all full-length POLDIP2 models.** (a) cartoon representation of Robetta models. Domain colouring as in Figures 1 and 2a, excepting N-terminal domain in blue cartoon representation (NTD) and modelled loops as dashed lines. C-term, C-terminus. (b) 100 ns molecular dynamics simulation of Robetta models (orientated with respect to respective model in panel (a), with colour scale and chain thickness representing RMSD.

FIGURE S5

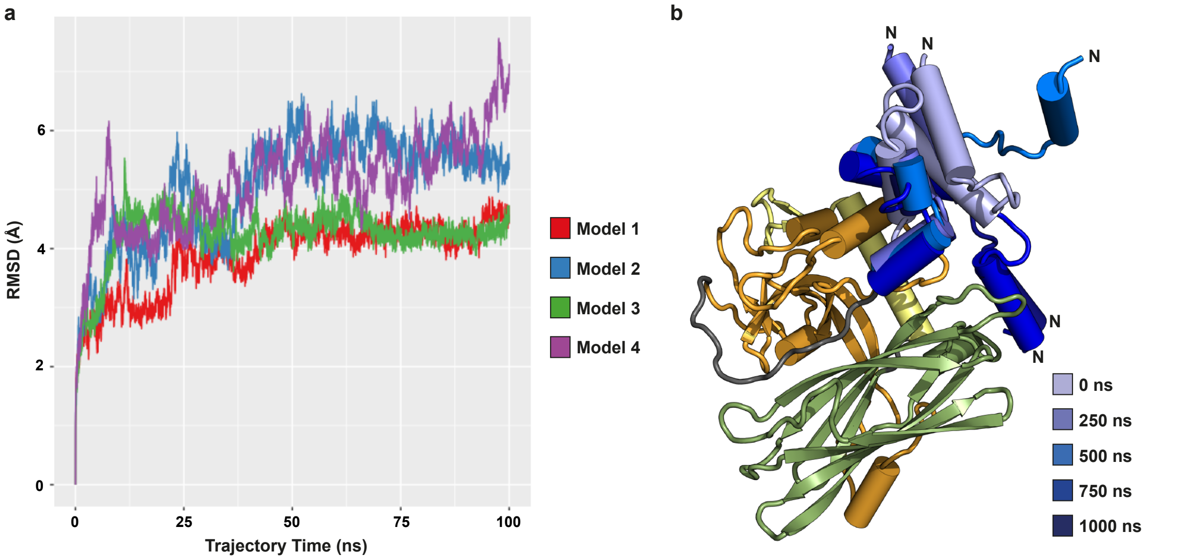


**Supplementary Figure S5. POLDIP2$^{FL}$ molecular dynamics time simulations.** (a) root mean square deviation (RMSD, deviation of model over time from reference position) simulation over 100 ns. (b) cartoon representation of NTD dynamics of POLDIP2$^{FL}$ model 4 1000 ns simulation. Core structure is model 4 at time 0 ns, with NTD structures superimposed from five time points (N, N-terminus).
